# Biophysical modeling for accurate T cell specificity prediction of viral and tumor antigens

**DOI:** 10.1101/2025.05.25.655924

**Authors:** Zahra S. Ghoreyshi, Noah Tubo, Luca Zammataro, Xizeng Mao, Ho Ngai, Duncheng Wang, Yibin Chen, Qiuming He, Eduardo Cisneros, Shoudan Liang, Priya J. Koppikar, Xingcheng Lin, Jeffrey J. Molldrem, Jason T. George

## Abstract

Accurate predictions of T cell receptor (TCR) specificity remain an important open problem in immunology, with broad implications for vaccine design, optimal immunotherapy, and improved management of autoimmune diseases. However, diversity in peptide antigens and TCR sequences at the level of individual patient repertoires remains a formidable computational challenge. Here, we develop a joint experimental and computational approach for predicting the antigen specificity of clinically-derived TCR sequences. Our model is trained on a combination of experimentally pre-identified and *in silico*-predicted TCR-pMHC structures using AlphaFold3. We apply our structural model in the clinical setting of hematopoietic stem cell transplant (HSCT) and demonstrate that our model is able to effectively discern the specificity of previously unseen donor and patient-derived TCR sequences against tumor associated and viral antigens. Model performance was further enhanced through the integration of sequence-based clustering and structurally diverse training templates. Our results highlight the predictive capabilities of structurally guided machine learning frameworks, trained on a minority test dataset, for antigen specificity prediction on unseen TCR sequences and their potential impact on a wide range of immunological applications.

## Introduction

The ability of T cell receptors (TCRs) to selectively recognize short antigenic peptides bound to major histocompatibility complex (MHC) molecules underpins a broad spectrum of adaptive immune responses in contexts such as infection, cancer, and autoimmunity. Accurate prediction of peptide recognition by diverse TCR repertoires remains a central challenge in computational immunology, with relevance for vaccine design, immune monitoring, and the development of targeted immunotherapies. Human TCR repertoires are remarkably diverse — each comprising an estimated 10^8^ unique clonotypes drawn from a theoretical space of up to 10^19^ — and also capable of self-non-self discrimination within an extensive (~ 10^13^) peptide landscape This vast combinatorial complexity therefore necessitates the development of robust computational frameworks to complement experimental strategies aimed at understanding repertoire-level TCR specificity.

Recent significant advancements have been made in understanding TCR recognition, driven by both the emergence of high-throughput experimental datasets (1, 2) and concurrent development of advanced computational frameworks that accurately capture TCR-pMHC interactions (3–13). Among these computational approaches, structurally informed biophysical models of the TCR-peptide interaction (RACER-m) have emerged as particularly effective frameworks to understand TCR specificity (12, 13). While these models have demonstrated reasonable accuracy in predicting specificity for a variety of publicly available datasets, their capacity to accurately predict antigen specificity for previously unseen TCR sequences remains unstudied.

In this study, we leverage the RACER-m framework to distinguish tumor-specific TCRs from those targeting viral epitopes. The clinical context of allogeneic hematopoietic stem cell transplantation (allo-HSCT) provides an ideal setting to evaluate and apply this strategy, as patient- and donor-derived expanded TCR repertoires are efficiently obtained and sequenced directly from peripheral blood or bone marrow samples. Following sequencing, TCR specificity was computationally predicted and subsequently validated experimentally through affinity-driven tetramer sorting, then evaluated iteratively. To address the limitation of sparse TCR training data in a discrete ligand space, we incorporated sequences obtained from HSCT-derived repertoires, experimentally validated TCR-pMHC complexes, and additional *in silico*-derived TCR-pMHC structural models generated using several recent computational approaches, including AlphaFold3 (14). Furthermore, we established peptide-specific binder distribution profiles to more accurately characterize antigen recognition. The integration structural features and sequence-based clustering via TCRdist (15, 16) significantly enhanced the model’s ability to resolve the specificity of previously unseen TCRs against viral and tumor antigens. Our specialized approach consistently achieved high predictive accuracy (>80%) across multiple TCR test cohorts. Collectively, our results illustrate the predictive power of structurally informed models. This joint experimental and modeling approach also establishes an iterative strategy for continuous model refinement through sequential incorporation of new structural and repertoire data as they become available, which we anticipate will be of broad use in diverse clinical contexts (13, 17–20).

## Materials and Methods

### Experimental Methods

#### Generation of peptide:HLA-I monomers and tetramers

Peptide-loaded biotinylated HLA monomers were produced via individual refolding reactions containing recombinant HLA-A^*\**^02:01, B2M, and a fixed peptides (21), or recombinant HLA-A^*\**^02:01, B2M, and a UV-exchangeable peptide (22). HLA monomers were sourced from the Baylor College of Medicine Tetramer Core and Biolegend. Peptides were synthesized using standard Fmoc-chemistry and purified to >90% via LC-MS (Genscript).

Peptide:HLA-I with the desired peptide specificities were generated via UV-mediated ligand exchange, as described elsewhere (22). Briefly, HLA monomer was diluted to 0.1 mg/ml in PBS and mixed 1:1 with 400uM target peptide in PBS. The mixture was transferred to a polypropylene 96-well plate and exposed to 365nm UV-C light for 30 min using a CL-3000 UV crosslinker (Analytik Jena), followed by overnight incubation at 4°C. The efficiency of UV-exchange reaction was assessed using the LEGEND MAX Flex-T Human Class I Peptide Exchange ELISA kit (Biolegend), which measures the association of biotinylated MHCI and B2M (Biolegend).

Peptide:HLA tetramers were produced by adding streptavidin-conjugates to biotinylated monomers in four serial additions, with 10-minute incubations between additions, such that the final stoichiometric ratio of streptavidin to peptide:MHCI was 1:4. Subsequently, any remaining unoccupied biotin-binding sites were quenched by the addition of 50nM soluble D-biotin. Fluorescent tetramers were made by conjugating HLA monomers with premium-grade phycoerythrin (PE) and/or allophycocyanin (APC) streptavidin conjugates (Invitrogen). DNA-barcoded tetramers were produced by linking monomers to TotalseqC-streptavidin conjugates (Biolegend). Conjugations were carried out on ice.

#### Tetramer staining and flow cytometry

TCRs were initially isolated based on their relative abundance in peripheral blood or bone marrow samples obtained from healthy donors. Cells were stained using tetramers specific for several antigens, including MART-1, FLU, and CMV-NLV, and subsequently enriched by magnetic selection to isolate tetramer-binding populations. The enriched cells then underwent approximately 10 days of *in vitro* expansion, using anti-CD3/CD28/CD2 stimulation supplemented with IL-2. After expansion, the cells were stained again with DNA-barcoded tetramers, each antigen specificity uniquely encoded by a distinct combination of two DNA barcodes. Tetramer-positive cells were pooled, sorted, and subsequently sequenced to identify paired TCR sequences along with their associated DNA barcodes. Cells were assigned antigen specificity based on robust binding to the correct DNA barcode combination at the single-cell level, without significant cross-reactivity. A TCR clonotype was defined as antigen-specific only if greater than 90% of cells sharing that clonotype demonstrated the same antigen specificity. These TCR sequences thus represent experimentally identified antigen-specific TCRs, rather than functionally confirmed receptors.

Cells were resuspended to 25M cells/ml in staining buffer (PBS + 2% FBS + 2mM EDTA) containing 50nM Dasatinib. Cells were stained on ice with 5ug/ml of fluorescent and/or DNA-barcoded peptide:MHCI tetramers for 30 minutes. Cells were washed, and subsequently stained with a cocktail of antibodies including CD8a-Buv395 (RPA-T8, BD), CD3e-BV421 (OKT3, Biolegend), Dump-FITC (CD4 (RPA-T4), CD19 (HIB19), CD14 (M5E2), CD16(3G8)), anti-APC (APC003, Biolegend), anti-PE (PE001, Biolegend). In some experiments anti-streptavidin-PE (3A20.2, Biolegend) was added. For endpoint analysis cells were run on a FACS Symphony A5 (BD) and analyzed using FlowJo.

For sequencing experiments, live, tetramer-positive, dump-negative CD8 T cells were sorted. Cells were immediately run using the 10X Genomics 5’ Single Cell Immune Profiling workflow, using the manufacturer’s instructions. Briefly, the sorted cells were loaded onto a lane of a Chromium Next GEM Chip G (10x Genomics) for GEM production using the Chromium controller. GEM-RT, cDNA amplification and the following library prep were using the Chromium Single Cell V(D)J Reagent Kits User Guide (v1.1 Chemistry) with Feature Barcoding technology for Cell Surface Protein (RevG) workflow (10x Genomics) as per 10x Genomics protocols.

#### Illumina sequencing

DNA libraries including Feature Barcode, TCR and GEX (gene expression) were quality controlled using TapeStation D5000 high sensitivity tape (Agilent). TapeStation bioanalyzer readout was also referenced for average DNA size calculation to determine the library loading for multiplexed sequencing. Concentration of library dsDNA was measured using a Qubit dsDNA Broad Range kit (Life Technologies, cat. Q32850). Libraries were sequenced on an Illumina NovaSeq 6000 S1 reagent kit v1.5 100 cycles (Illumina 20028319) using sequencing cycles configuration of 26+8+0+91 (Read 1 + i7 Index + i5 Index + Read 2) for combined assessment of feature barcode library, TCR and GEX (RNA seq) information. Sequencing depth is 7,500 read pairs per cell for feature barcode library, 7,500 read pairs per cell for TCR library, and 20,000 read pairs for GEX library.

### Model Development

To predict binding affinity between TCR-peptide pairs, we specialized our previously developed RACER-m model, an energy-based biophysical framework designed to quantify TCR–pMHC interactions (12, 13). RACER-m focuses on the CDR3*α* and CDR3*β* regions as the primary mediators of context-specific TCR-peptide binding (23). We retrained and optimized RACER-m model using 66 experimentally derived TCR–pMHC structures as the initial core structural dataset. TCR sequence used in training included 97 binder TCR–peptide pairs from the ATLAS database (24), as well as additional TCR–pMHC complexes sourced from MDACC and experimentally confirmed as binders (see Table 1). For each strong-binding TCR–peptide pair, we generated an additional set of 1,000 random TCR sequences to serve as non-binding examples during training. Threading these TCR–peptide pairs through candidate structural templates selected based on sequence similarity enabled the development of an optimized energy matrix for accurately predicting TCR–peptide binding affinities in this clinical context.

**Table 1.**
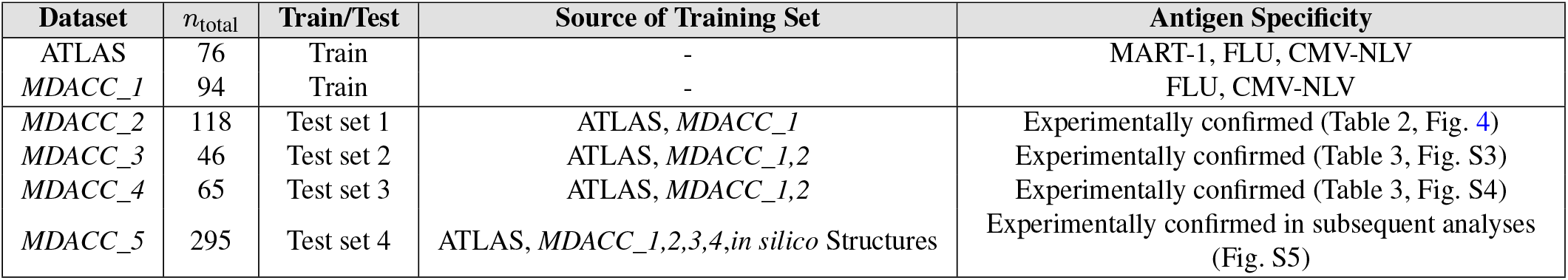
Summary of datasets used in this study. The table lists the total number of TCR cases (*n*_total_) in each dataset and specifies their role as either training or test sets. The ATLAS dataset and *MDACC_1* comprise the primary training datasets, containing known antigen specificities for MART-1, FLU, and CMV-NLV. The MDACC indices (*MDACC_2* to *MDACC_5*) represent distinct donor and patient-derived TCR cohorts from MD Anderson Cancer Center, each sequentially evaluated as test sets. Specific antigen distributions for these test datasets were experimentally validated and are comprehensively detailed in Tables 2 and 3, and further analyses.

| Dataset | $n_{\text{total}}$ | Train/Test | Source of Training Set | Antigen Specificity |
| --- | --- | --- | --- | --- |
| ATLAS | 76 | Train | - | MART-1, FLU, CMV-NLV |
| MDACC_1 | 94 | Train | - | FLU, CMV-NLV |
| MDACC_2 | 118 | Test set 1 | ATLAS, MDACC_1 | Experimentally confirmed (Table 2, Fig. 4) |
| MDACC_3 | 46 | Test set 2 | ATLAS, MDACC_1,2 | Experimentally confirmed (Table 3, Fig. S3) |
| MDACC_4 | 65 | Test set 3 | ATLAS, MDACC_1,2 | Experimentally confirmed (Table 3, Fig. S4) |
| MDACC_5 | 295 | Test set 4 | ATLAS, MDACC_1,2,3,4,in silico Structures | Experimentally confirmed in subsequent analyses (Fig. S5) |

#### Model training and optimization

Following the procedure outlined in (12, 13), we trained the energy model to maximize the energy gap, *δE/*Δ*E*, between binders and non-binders. For each binder, the average binding energy, ⟨*E*_binder_⟩, was computed, along with the mean energy of its corresponding non-binders, ⟨*E*_non-binder_⟩, and the standard deviation Δ*E* of the non-binder energies. This approach maximizes the ratio *δE/*Δ*E*, where *δE* = ⟨*E*_non-binder_⟩− ⟨*E*_binder_⟩, measures the separation between binders and non-binders. RACER-m achieves this optimization by maximizing the ratio *δE/*Δ*E* using an adapted AWSEM force field (25), a residue-level protein force field widely used for investigating protein folding and interactions (1, 25). We utilized the protein-protein interaction component of the force field to compute contact energies specifically at the TCR-peptide interface. Specifically, *Cβ* atoms were employed for most residues, except for glycine, where *Cα* atoms were used. This calculation involves interaction weights for residue pairs and distance-based potentials to compute binding energies. The binding energy is given by:

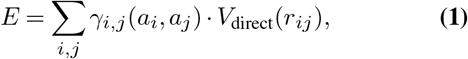

where *γ*_*i,j*_(*a*_*i*_, *a*_*j*_) represents interaction weights for residue pairs *i* and *j* with amino acid types *a*_*i*_ and *a*_*j*_, respectively, and *V*_direct_(*r*_*ij*_) denotes the direct interaction potential as a function of the inter-residue distance *r*_*ij*_.

For unknown TCRs, RACER-m applies template-based modeling with MODELLER (26), matching test sequences to template structures based on composite similarity scores across CDR3*α*, CDR3*β*, and peptide regions (Figure 1).

**Fig. 1.**
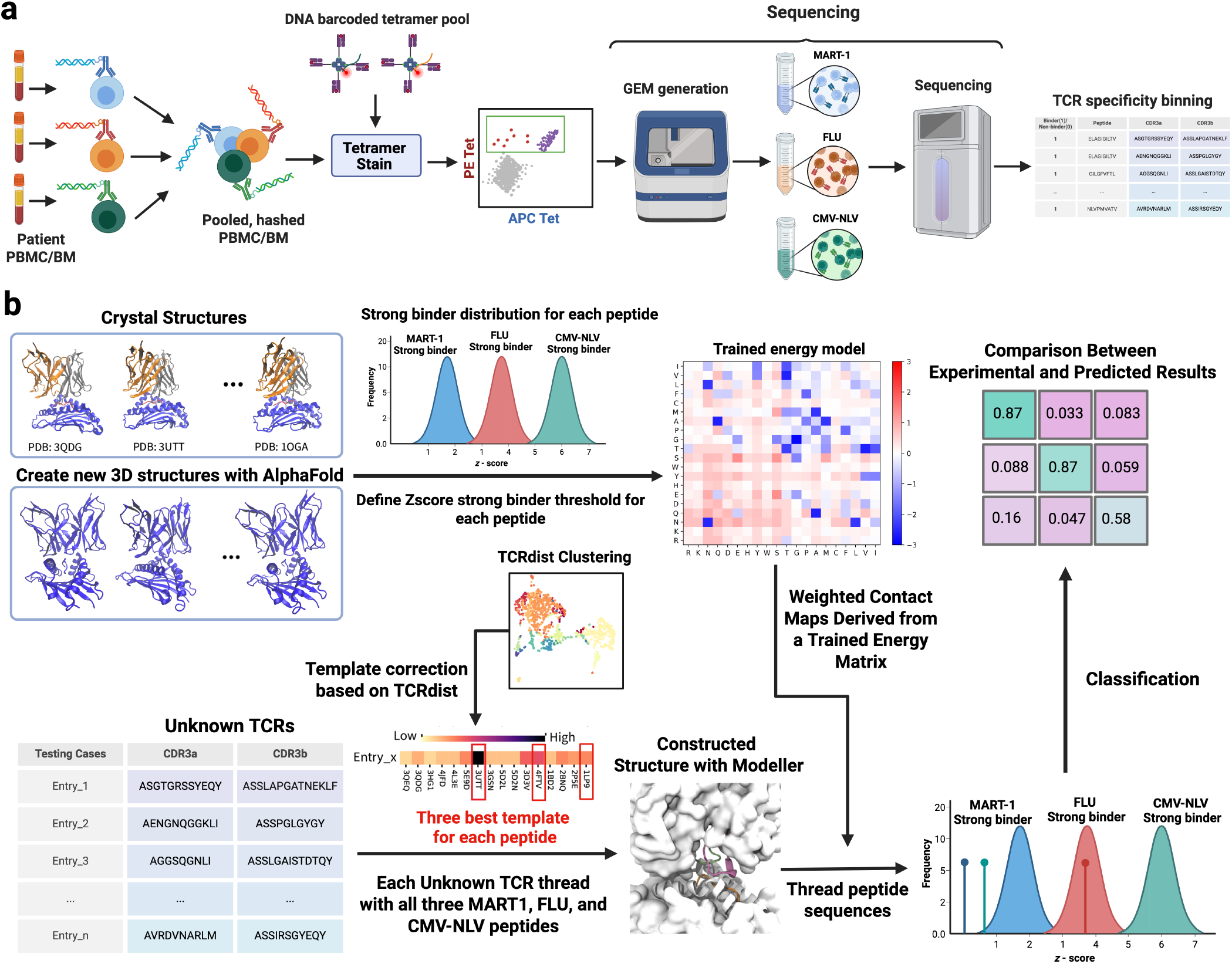
Schematic representation of the workflow. (**a**) Experimental setup: Patient-derived and healthy donor peripheral blood mononuclear cell (PBMC) or bone marrow (BM) samples are labeled with sample-specific hashtags, pooled, and stained with a DNA-barcoded tetramer library, where each tetramer is uniquely encoded by a specific DNA barcode. Tetramer-binding (PE and/or APC) cells are sorted and sequenced to determine both tetramer identity and the paired TCR*α/β* sequences. (**b**) Training and testing pipeline: the training phase utilizes crystal structures and synthetic *in silico* data (e.g., AlphaFold-derived TCR–pMHC complexes) to build or refine the predictive model. The testing phase classifies unknown TCRs, predicting their recognition of one of several candidate peptides (e.g., MART-1, FLU, CMV-NLV).

#### TCR-pMHC structural modeling

Since RACER-m evaluates binding energies based on contact interactions between peptides and TCRs, it relies on the availability of 3D structures for TCR-pMHC complexes. Obtaining this information for every test case is impractical, particularly when the relevant task involves characterizing specificity for subsets of the T cell repertoire that are distinct from training sequences. To address this limitation, RACER-m (13) utilizes `MODELLER` (26) to build 3D models from sequence data, which we adopt in this study.

For each TCR-pMHC test pair, RACER-m computed Hamming distances separately for the peptide, CDR3*α*, and CDR3*β* sequences relative to each of the 66 template structures. Sequence similarity scores were calculated by counting the number of matching amino acids between the target and template sequences. A composite similarity score was then obtained by summing the similarity scores of the CDR3 regions and multiplying by the peptide similarity score. The template with the highest composite similarity score was selected for structure generation using MODELLER (see Figure 1-b).

#### Similarity-based template correction

To improve the accuracy of template selection in the RACER-m model (12), we employed a two-part correction strategy based on structural and sequence similarity measures. The initial RACER-m model used a training set of 66 crystal structures and organized them into clusters using the mutual Q similarity metric (27, 28). Mutual Q quantifies structural similarity between 3D configurations, allowing for global comparison of two protein structures. This metric, calculated as:

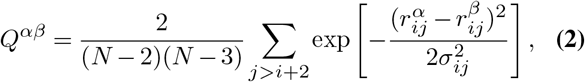

where *N* represents the total number of residues,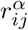 and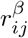denote the distances between residues *i* and *j* in configurations *α* and *β*, respectively, and *σ*_*ij*_ accounts for fluctuations. This method groups structures into clusters based on global structural similarities. While effective, this approach has limitations, particularly in cases where possible structural variations within diverse TCR sequences affect binding affinity.

To address these limitations, we incorporated TCRdist (15, 16), a sequence-based clustering method that refines template grouping using CDR3*α* and CDR3*β* gene sequences. TCRdist computes pairwise distances between TCRs by comparing concatenated CDR loop sequences using a similarity-weighted Hamming distance. This calculation incorporates amino acid substitution penalties from the BLOSUM62 matrix (29) and applies gap penalties to account for length variation, with greater weight assigned to the CDR3 region due to its critical role in antigen specificity. This weighted metric emphasizes sequence features that are functionally important for pMHC recognition, enabling more accurate clustering of TCRs based on their positional and biochemical similarity.

#### In silico structure generation

In the RACER-m model, the relative scarcity of crystal structures was previously shown to limit accurate predictions on structurally disparate TCR-peptide systems (13). We will show that this limitation was especially pronounced for patients with diverse TCR clusters recognizing a specific peptide, such as MART-1, where the existing crystal structures were insufficient for accurate predictions. To address this limitation, we utilized AlphaFold3 (14) to generate additional 3D structures from a minority (~10%) of test cases, incorporating these into the training set. Specifically, from the full set 𝒟 = {*d*_1_, *d*_2_, …, *d*_*N*_} of *N* TCRs obtained and sequenced in a given specificity experiment, we randomly select a minority fraction (*N/*10) for specificity determination and template construction. Following complete experimental validation of our results, we then iterated this same procedure *K* = 6 times to establish confidence intervals for our results. This approach allowed us to assess the reliability of the generated subsets and identify the best set of structures to augment the training data, thereby improving the model’s predictive performance.

In a few cases where we observed overlap in TCR primary sequences between training and test sets, we used this as an opportunity to generate and evaluate new *in silico*-derived structures for the overlapping TCRs that lacked structural information using AlphaFold3. When such sequences appeared in the randomly selected training subset, we replaced their original entries with the newly modeled structures. Predictive performance for these updated models was assessed by evaluating their ability to distinguish binding versus non-binding cases using ROC curve analysis, repeated over multiple randomized training and test splits to ensure robustness. To comprehensively evaluate any potential effect of utilizing different methodology for *in silico* template creation on predictive performance, we repeated the above procedure using AlphaFold Multimer (30), along with more recent structure prediction methods, including Boltz-1 (31), which leverage large-scale transformer-based architectures and diffusion models designed for biomolecular complex prediction. We then evaluated these models as done above for AlphaFold3.

To generate new structures for the excluded cases, we required detailed sequence information, including pMHC sequences and TCR variable regions (*V*_*α*_ and *V*_*β*_). For the testing cases, sequence data were available only for CDR3*α*/*β* and the peptide, with the MHC being identical across all cases (HLA-A^*\**^02:01). To obtain the remaining V*α*/*β* sequences, we referred to the best template identified by RACER-m during the primary selection step. These sequences were then used to construct new 3D structures, which were incorporated into the training set to enhance model applicability. Similarly, for non-CDR regions, we selected consensus sequences derived from curated TCR databases and validated structural templates to ensure proper folding and structural stability. The complete TCR sequences—including both CDR and non-CDR regions—along with the pMHC sequences, were then used to generate new 3D structures.

#### TCR-pMHC distribution percentile calculation

The recognition task in this study focuses on identifying T cell receptors (TCRs) as binders to a single antigen target by learning distinctive biophysical features from training distributions of binders and non-binders. We implemented a new classification method that improved upon our previous approach (RACER-m) (13) that originally classified TCR-pMHC pairs as binders based on a global *Z*-score cutoff. Using RACER-m, we generated *Z*-score distributions across our training set for three antigens: MART-1, FLU, and CMV-NLV. For an unknown TCR, we computed *Z*-scores for each of these antigen pairs and determined the percentile ranking of each *Z*-score within its respective distribution. The percentile rank serves as a measure of binding potential for each peptide based on its *Z*-score (Figure 2A). In this approach, the peptide having the highest percentile ranking is identified as the strongest binder. When the *Z*-score exceeds the mean, a higher percentile indicates a stronger binding interaction and reinforces its classification as a binder. Conversely, when the *Z*-score falls below the mean, larger percentile scores correspond to values closer to the mean, signifying that among the available options, this peptide exhibits the highest relative binding potential. This formulation ensures a consistent and systematic selection of the most probable binder across all scenarios.

**Fig. 2.**
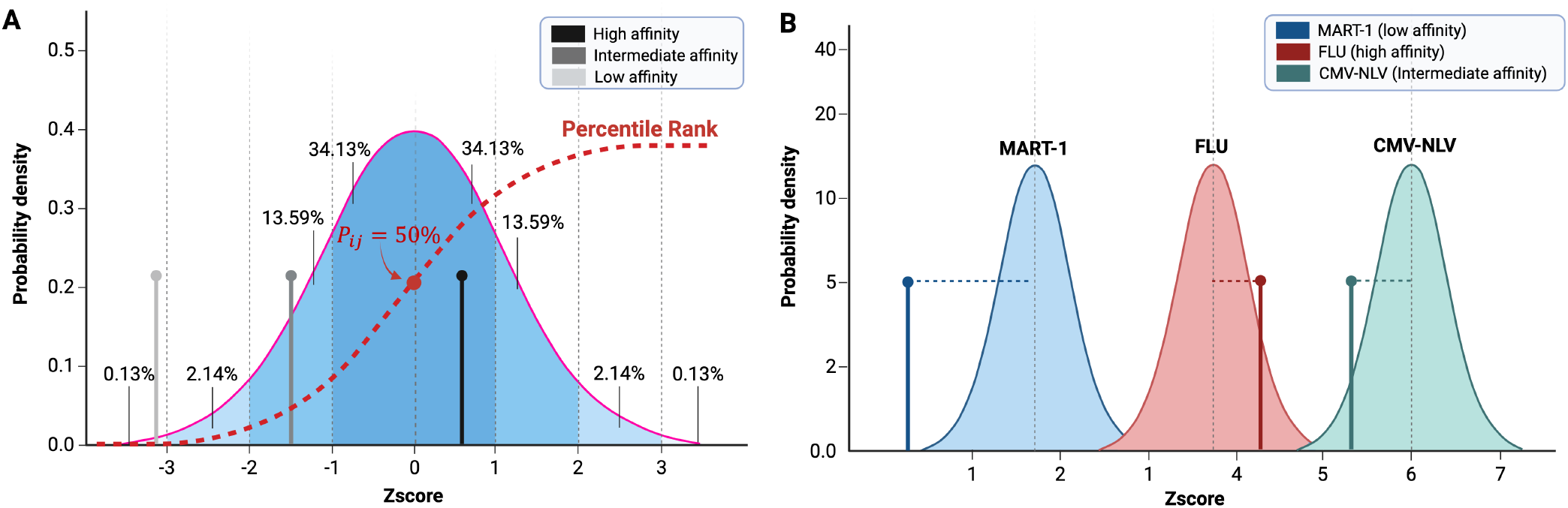
Z-score binding criteria. (a) Binding confidence is determined by *Z*-score relative to the distribution of known binders. Confident (dark gray), intermediate confidence (gray), and low confidence (light gray) is assigned to determine binding and non-binding TCR-peptide pairs. (b) TCR Selection criteria for determining binding/non-binding is determined by identifying the peptide exhibiting maximal z-score.

The binding strength of each peptide is expressed as a percentile rank (*P*_*ij*_), which represents the percentile rank of TCR *i* amongst the distribution of known binder TCRs 𝒯_*j*_ for peptide *j* out of a collection of peptides 𝒫. These distributions are characterized by their respective means (*µ*_*j*_) and standard deviations (*σ*_*j*_), derived from the training data. The percentile for each peptide is calculated using the cumulative distribution function (CDF) of the normal distribution:

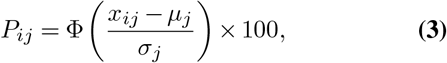

where *x*_*ij*_ is the observed binding strength for TCR *i* in the set 𝒯_*j*_ of TCRs specific for peptide *j*, and Φ is the CDF of the standard normal distribution. The percentile provides an interpretable measure of the binding strength relative to the distribution, enabling meaningful comparisons among the peptides. The specificity of TCR *i, S*_*i*_, can be identified by maximizing over the available TCR percentiles for each peptide:

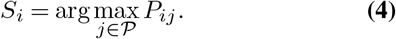

This selection criterion associates each TCR with the peptide showing the maximal percentile binding affinity (Figure 2b). This approach evaluates the binding strength of each TCR test case against the three peptides (MART-1, FLU, and CMV-NLV) from which sensitivity, specificity, and diagnostic accuracy are quantified. The selection criteria are designed to ensure that TCRs are classified as specific to exactly one peptide from this list.

## Results

### Optimized structural biophysical modeling successfully recovers TCR specificity

We specialized the original RACER-m framework with the goal of generating a model capable of resolving viral (CMV-NLV, FLU) and cancer (MART-1) epitope-specific TCR sequences (13). The publicly available TCRs in the original model construction exhibited significant class imbalance, with a higher representation of MART-1-specific TCRs (*n*_MART-1_ = 53) compared to viral (*n*_FLU_ = 11; *n*_CMV-NLV_ = 12) examples. To address this imbalance and also include clinically derived cancer- and viral-specific TCRs, we experimentally assessed specificity for a variety of TCRs obtained from HSCT donors and patients. In particular, the ATLAS database (24) was augmented by the addition of 20 FLU-specific and 74 CMV-NLV-specific experimentally confirmed TCRs (‘*MDACC_1*’; Table 1) (see Methods).

Following training and optimization, we analyzed the distributions of normalized (Z-score) affinity values of TCRs specific to each of MART-1, FLU, and CMV-NLV (Figure 3). To refine the selection of structural templates, we quantified the structural similarity of within-family TCRs by applying the mutual Q similarity approach (27, 28) (see Materials and Methods). We then excluded templates that were significantly distant from others within their respective specificity groups. For example, PDB ID 5HNT, with its relatively low (3.2 Å) resolution, exhibited minimal similarity with all other MART-1-specific TCR-pMHC structures and was thus excluded. Similarly, additional templates for MART-1, FLU, and CMV-NLV were identified as outliers and removed based on mutual Q similarity (Table S1 in the Supplementary Information). Template exclusion was performed because such outliers lead to an inaccurate estimation of percentile assignment, and their exclusion was shown to improve percentile mappings (Figure S1).

**Fig. 3.**
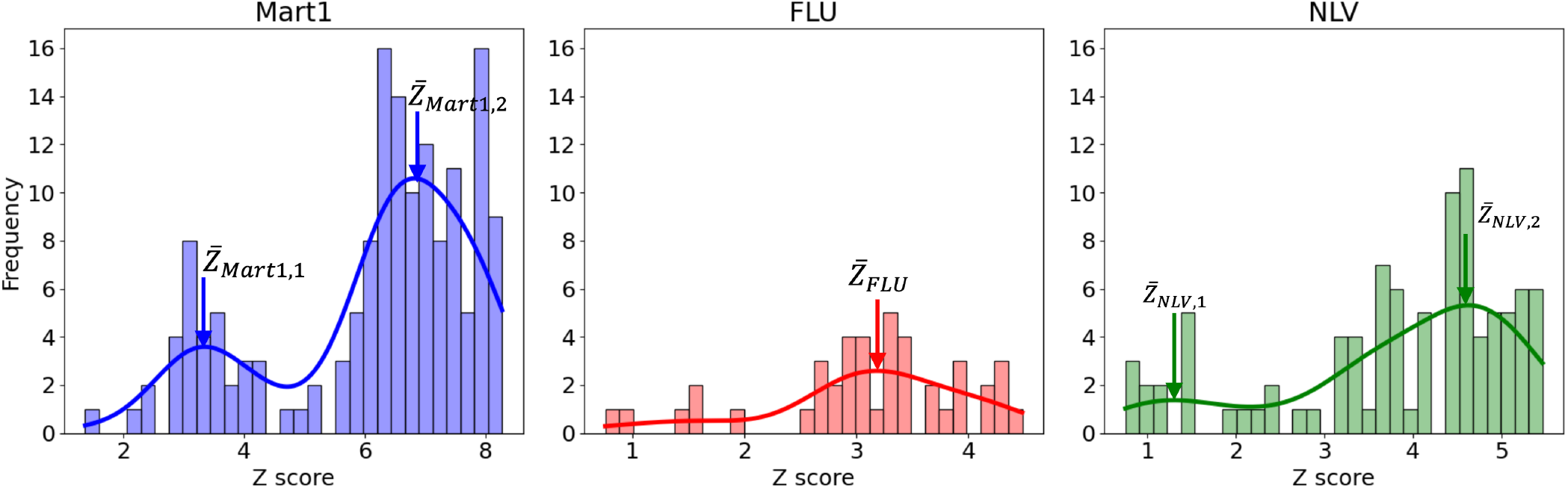
Distribution of *Z*-scores for MART-1 (blue), FLU (red), and CMV-NLV (green) in Training Set 1, which includes the ATLAS and *MDACC_1*. The MART-1 and CMV-NLV distributions exhibit two distinct peaks, indicating the presence of two different structural conformations, whereas the FLU distribution shows a single peak. These *Z*-score distributions were obtained after refining the structural template library using mutual Q similarity metrics (see SI sec. S1, Table S1, and figure S1) to select the optimal templates for binding energy calculations.

Repeating the analysis with inaccurate templates elsewhere yielded two distinct clusters for *MART-1*, one cluster for *FLU*, and two clusters for *CMV-NLV*, as visualized by hierarchical clustering (Figure S1). Consequently, we established distinct statistical tests corresponding to each structural cluster based on the identified Z-score distributions. *MART-1* cases exhibited two empirical distributions with means of *µ*_MART-1,1_ = 3.2 and *µ*_MART-1,2_ = 6.8. *CMV-NLV* cases also displayed two distinct distributions with means of *µ*_CMV-NLV,1_ = 1.3 and *µ*_CMV-NLV,2_ = 4.6. In contrast, *FLU* cases were characterized by a single distribution with a mean of *µ*_FLU_ = 3.2. All distributions were normalized to a standard deviation of *σ* = 1 (Figure 3). We integrated these distributions into our model by assigning each test case to the appropriate structural cluster using sequence-based similarity to the closest template. Because each structural cluster corresponded uniquely to one statistical distribution, we then performed the respective statistical test by calculating the within-cluster percentile rank. The resulting percentile rank thus quantified the relative binding strength of the TCR to each pMHC complex for the three target peptides analyzed.

To evaluate optimized model performance, we predicted the specificity of 118 HLA-A^*\**^02-restricted TCRs from a independent PBMC and BM-derived TCR cohort (*MDACC_2*; Table 1). The ROC curves for *MDACC_2* are summarized in Figure 4, while Table 2 reports the sensitivity, specificity, and diagnostic accuracy. The model demonstrated strong predictive performance across all three peptides, with the highest accuracy for *CMV-NLV* (0.91), followed by *FLU* (0.90) and *MART-1* (0.85). Moreover, this refined template selection using mutual Q similarity contributed to improving classification performance, particularly by ensuring that the structural representations used for *MDACC_2* cases were well aligned with the training distributions (Figure S1).

**Table 2.** Specificity, Sensitivity, and Accuracy for MART-1, FLU, and CMV-NLV using *MDACC_2*.

| Peptide | Specificity | Sensitivity | Accuracy |
| --- | --- | --- | --- |
| MART-1 ($n = 97$) | 0.92 | 0.85 | 0.85 |
| FLU ($n = 9$) | 0.95 | 0.88 | 0.9 |
| CMV-NLV ($n = 12$) | 0.96 | 0.89 | 0.91 |

**Fig. 4.**
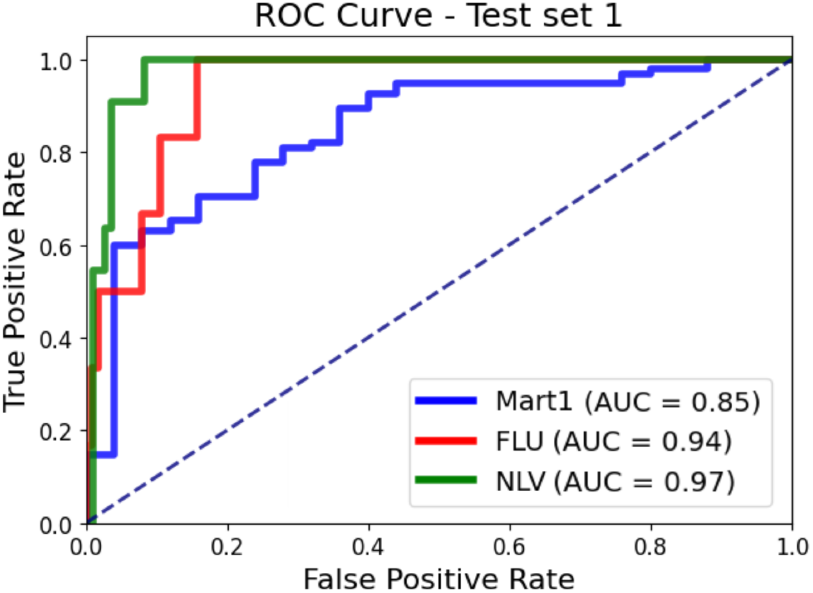
Receiver Operating Characteristic (ROC) curve results for test set 1, using the mutual Q method as the criterion for clustering TCRs specific to MART-1, FLU, and CMV-NLV.

### Sequence similarity-based clustering enables appropriate template selection in testing

Given our model’s initial predictive performance, we next aimed to develop an update policy whereby the specificities of previously obtained donor-derived TCR sequences, once experimentally confirmed, can be incorporated into model training. We thus performed a series of repeated tests of our modeling framework on additional independent datasets. The prior training and experimentally confirmed test data were incorporated into an updated training dataset (comprised of ATLAS, *MDACC_1*, and *MDACC_2* sequences). Two additional independent test datasets (*MDACC_3* and *MDACC_4*) from PBMC and BM-derived TCR cohorts were then used to assess the model’s ability to generalize across diverse TCR repertoires. In all subsequent test cases, any duplicate TCR sequences were removed from predictions.

In our initial test (*MDACC_2*), a structure-based template selection approach effectively mapped test sequences with an appropriate structure. However, when we applied this same method to additional unseen TCR cohorts (*MDACC_3* and *MDACC_4*), our initial approach failed to maintain predictive accuracy (Figures S3b and S4b). A careful examination of the inverse mapping (*f* ^*−*1^ : *Y*→ *X*) from the test TCR sequences *Y* to the set of structural templates available in training *X* revealed sensitivity issues, wherein test sequences mapped to multiple templates in some cases, while in others, similar test sequences were mapped to disparate structures for threading. As a result, we turned to TCRdist (15, 16), a previously established sequence-based clustering method that leverages detailed sequence similarity metrics to partition TCR sequences into distinct clusters. Specifically, we defined an optimized clustering threshold by selecting the largest TCRdist radius around an antigen-associated (centroid) TCR for which fewer than one in one million (10^*−*6^) background TCRs are expected to fall. This threshold was determined by constructing empirical cumulative distribution functions for each candidate centroid TCR using both antigen-associated sequences and a specifically constructed background repertoire comprising V–J gene-matched synthetic sequences and cord blood TCRs. By selecting this threshold, we maximized the capture of closely related antigen-associated TCRs while minimizing the inclusion of nonspecific, background TCR sequences. Although this approach inevitably groups together sequences with subtle structural variations, it strikes a balance between sensitivity (capturing relevant TCR sequences) and specificity (excluding unrelated background sequences).

We performed TCRdist clustering on the updated training set (ATLAS, *MDACC_1*, and *MDACC_2*) using CDR3*α/β* sequences and genetic features (J-gene and V-gene), available from the Immune Epitope Database (IEDB) (32) and germline sequenced MDACC datasets (excluding ATLAS sequences lacking J-gene and V-gene information). This clustering was then used to restrict candidate structural templates to those representing multiple closely related TCR sequences. Applying TCRdist to *MDACC_3* and *MDACC_4* refined template selection by filtering out singleton clusters. This method improved template selection for MART-1, FLU, and CMV-NLV, ensuring a prioritized list of structural templates accurately reflected the diversity of TCR sequences (Figure S2).

Using the updated TCRdist criteria for template matching, we then retrained our model and tested using the new test datasets. These predictions were then compared against experimentally determined ground-truth specificity for each TCR, which was also used to assess clustering accuracy within the TCRdist framework. The ROC analysis demonstrated that the model effectively distinguished binder TCRs across different epitope groups. The AUC values for MART-1 were 0.89 and 0.83 for *MDACC_3* and *MDACC_4*, respectively. For FLU, the corresponding values were 0.73 and 0.68, respectively, while for CMV-NLV, the model achieved AUC scores of 0.93 and 0.73, respectively. Notably, the limited number of FLU cases in *MDACC_3* and *MDACC_4* (only two cases each) may have influenced the lower reliability of the FLU predictions in these datasets. Figure 5a and Figure 5c present the ROC curves for *MDACC_3* and *MDACC_4*, while Figures 5b and 5d illustrate the corresponding TCRdist-based clustering results, with specificities indicated following experimental confirmation. In these clustering representations, clusters containing singleton TCRs were filtered for visualization purposes but remained included in the overall analysis. Moreover, these clustering results confirm that the structural families present in the training set (Figure S2) adequately cover the TCR diversity observed in *MDACC_3* and *MDACC_4*.

**Fig. 5.**
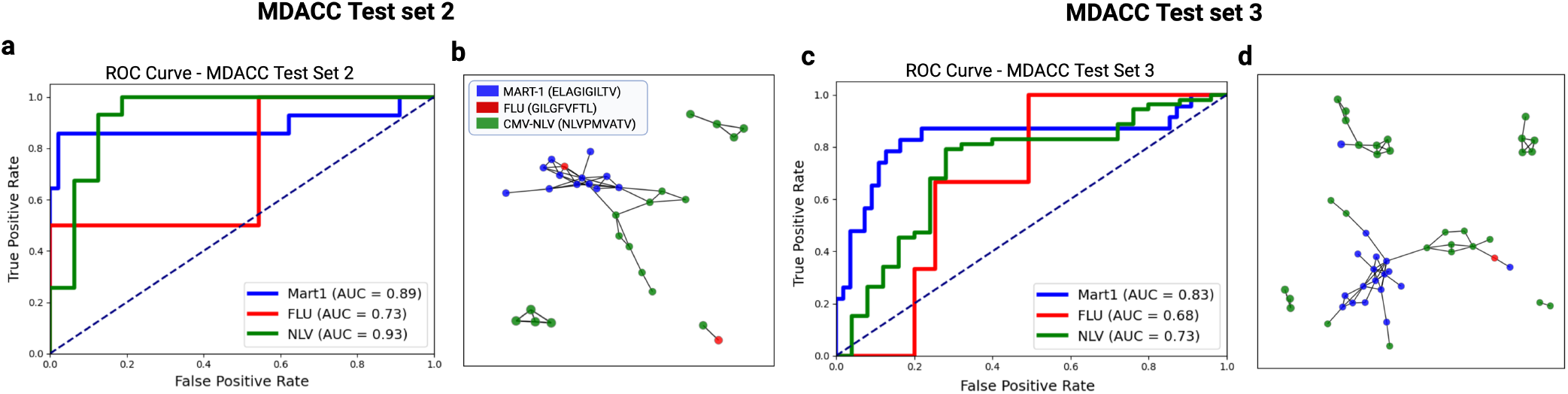
TCRdist-based analysis for *MDACC_3* and *MDACC_4*. (a, c)ROC curves demonstrating model performance in distinguishing binder TCRs for *MDACC_3* and *MDACC_4*, respectively. (b, d)Corresponding TCRdist-based clustering results, illustrating the sequence-based grouping of TCRs and confirming that the structural families in the training set (Figure S2) adequately cover the diversity observed in these datasets.

We also compared these results with an alternative classification strategy that focused solely on utilizing sequence-based classification of test TCRs based on TCRdist alone, thereby foregoing any structural modeling. While TCRdist provides a powerful clustering method based on sequence similarity, it alone is insufficient for accurate classification, there are numerous misclassifications among MART-1, FLU, and CMV-NLV peptides when relying solely on TCRdist clustering (Figure 5b,d). Precision (TP/(TP+FP)), Recall (TP/(TP+FN)), and Accuracy ((TP+TN)/(TP+TN+FP+FN)) were calculated for both methods (summarized in Table 3). Our results demonstrate that structural modeling combined with TCRDist-based template selection consistently achieves higher classification performance.

**Table 3.** Precision, Recall, and Accuracy for MART-1, FLU, and CMV-NLV using TCRdist and RACER-m methods on Test Set 2 and Test Set 3.

| Test Sets | Peptides | Precision |  | Recall |  | Accuracy |  |
| --- | --- | --- | --- | --- | --- | --- | --- |
|  |  | TCRdist | RACER+TCRdist | TCRdist | RACER+TCRdist | TCRdist | RACER+TCRdist |
| Test Set 2 | MART-1 ($n = 13$) | 0.55 | 1 | 0.84 | 0.76 | 0.71 | 0.93 |
| | FLU ($n = 2$) | 0 | 0.4 | 0 | 1 | 0.56 | 0.89 |
| | CMV-NLV ($n = 31$) | 0.62 | 0.93 | 0.7 | 0.87 | 0.52 | 0.84 |
| Test Set 3 | MART-1 ($n = 20$) | 0.44 | 0.71 | 0.84 | 0.75 | 0.63 | 0.72 |
| | FLU ($n = 2$) | 0.034 | 0 | 0.5 | 0 | 0.98 | 0.88 |
| | CMV-NLV ($n = 43$) | 0.96 | 0.78 | 0.74 | 0.7 | 0.78 | 0.72 |

### Incorporation of in silico-derived TCR-pMHC structures generalizes model predictions to diverse and unseen TCR test sequences

We further evaluated our approach using a final test dataset, comprising more patienttional templates would lead to improved model predictions.

To assess this possibility, we identified a small (n=15; ~5%) subset of maximally informative TCRdist-clustered test TCR sequences for the creation of new structural templates. Specifically, we selected TCRs from clusters containing four or more members and deliberately excluded those from highly dense ones (particularly the two central clusters in Figure S6). Our rationale for this exclusion was based on the observation that densely populated clusters typically reflect sequences with limited variability and conserved, canonical motifs that are shared across individuals. These high similar TCR sequences tend to have conserved structural conformations, and as such, their structural space is likely well-represented by existing crystal structures. By excluding these clusters from our selection, our aim was to focus on less populated, more heterogeneous cases that may harbor under-explored structural configurations (10, 15).

Once identified, each case was *a priori* experimentally tested to confirm peptide specificity and subsequently excluded from predictive assessment. We then used these confirmed TCR-pMHC systems to create additional *in silico*-derived structures using AlphaFold3 (14). In each instance, sequence data were available only for CDR3*α*/*β* and the corresponding peptide; accordingly, we incorporated the remaining V*α*/*β* sequences from the closest available RACER-m-identified crystal structure along with HLA-A^*\**^02:01 to generate synthetic TCR-pMHC structures for these selected cases (Figure 6). Figure 7a illustrates the selection process, where gray nodes represent *MDACC_5* test TCRs, and pink nodes indicate the chosen cases for *in silico* structure generation and subsequent inclusion in the training set.

**Fig. 6.**
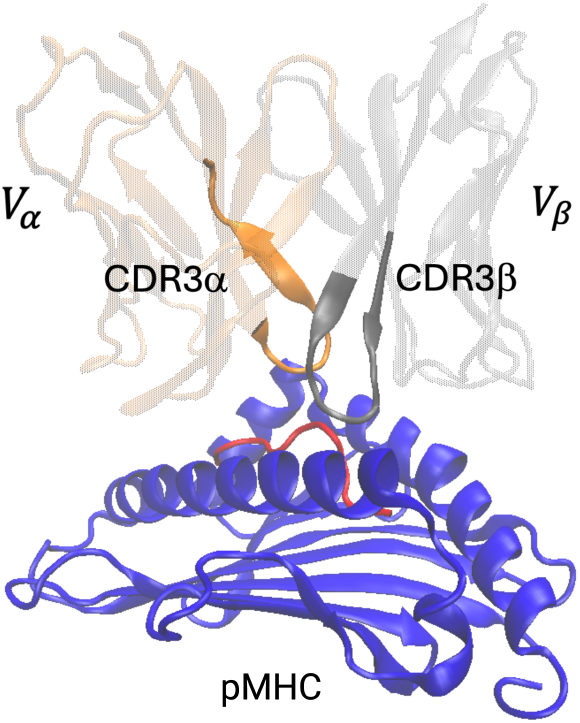
Structure generated by AlphaFold. The CDR3*α*/*β*, peptide, and MHC (colored regions) represent sequence information derived from the MDACC datasets, whereas the variable regions (*V*_*α*_ and *V*_*β*_), excluding CDR3*α*/*β*, are based on sequence information from the best template selected by RACER-m for each respective MDACC case.

**Fig. 7.**
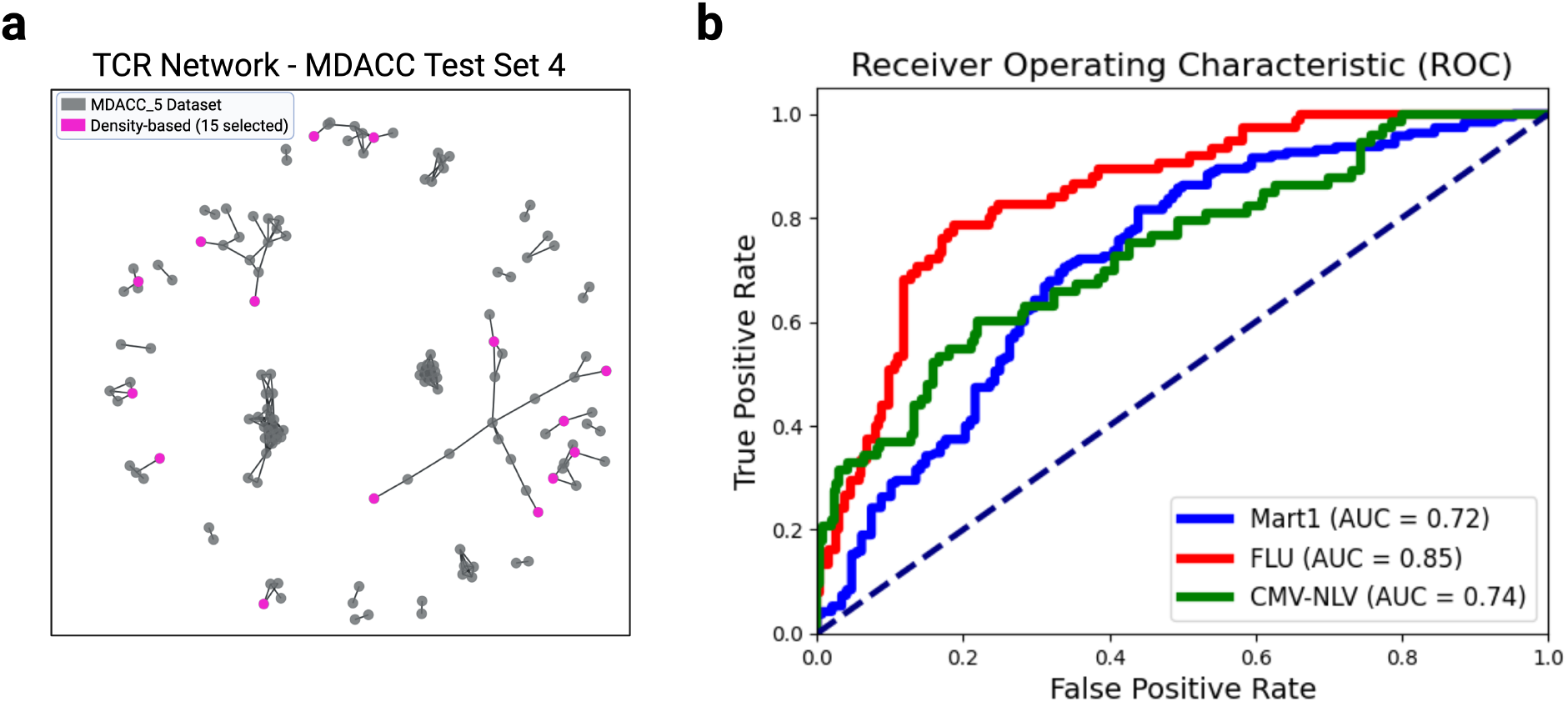
Comparison of the refined fifteen-case selection approach. (a) TCR clustering of the *MDACC_5* dataset, where gray nodes represent all TCRs, and pink nodes indicate the selected cases used for *in silico* structure generation and incorporation into the training set. (b) ROC curve showing the performance of the selected cases compared to the thirty-case random selection.

The ROC curve in Figure 7b illustrates the performance of this selection procedure, yielding ROC-AUC values of 0.72, 0.85, and 0.74 for MART-1, FLU, and CMV-NLV, respectively. A repeat attempt to create these predictions in the absence of new structures resulted in significant reductions in overall predictive performance (Figure S5), thereby quantifying the added benefit of utilizing an update strategy requiring minimal additional TCR information.

The above results were also compared with two alternative approaches. In the first alternative approach, a randomized procedure was utilized in place of the original density-based method to identify a subset (10%) of TCRs from the *MDACC_5* test set for structure generation. This was followed by predictive assessment on the 90% hold-out cases as a consistency check. We then retrained the RACER-m model using the same training procedure as before while incorporating the new structural templates. The optimized model was subsequently evaluated on each of the test datasets. This strategy effectively resolved duplication between training and testing sets while enriching the training data with structurally informative cases. This random selection was repeated over six iterations to establish confidence intervals on the ROC curves derived from randomized template selection (Figure 8a). Our repeat analysis following the generation of new structures for overlapping TCR sequences between training and test sets resulted in notable improvements in predictive performance. The ROC values across the different random subsets ranged from 0.69 to 0.76 for MART-1, 0.72 to 0.86 for FLU, and 0.64 to 0.74 for CMV-NLV (best and worst cases in Figure S7c and Figure S7d, respectively), all of which were comparable to the performance of the original density-dependent selection criterion.

**Fig. 8.**
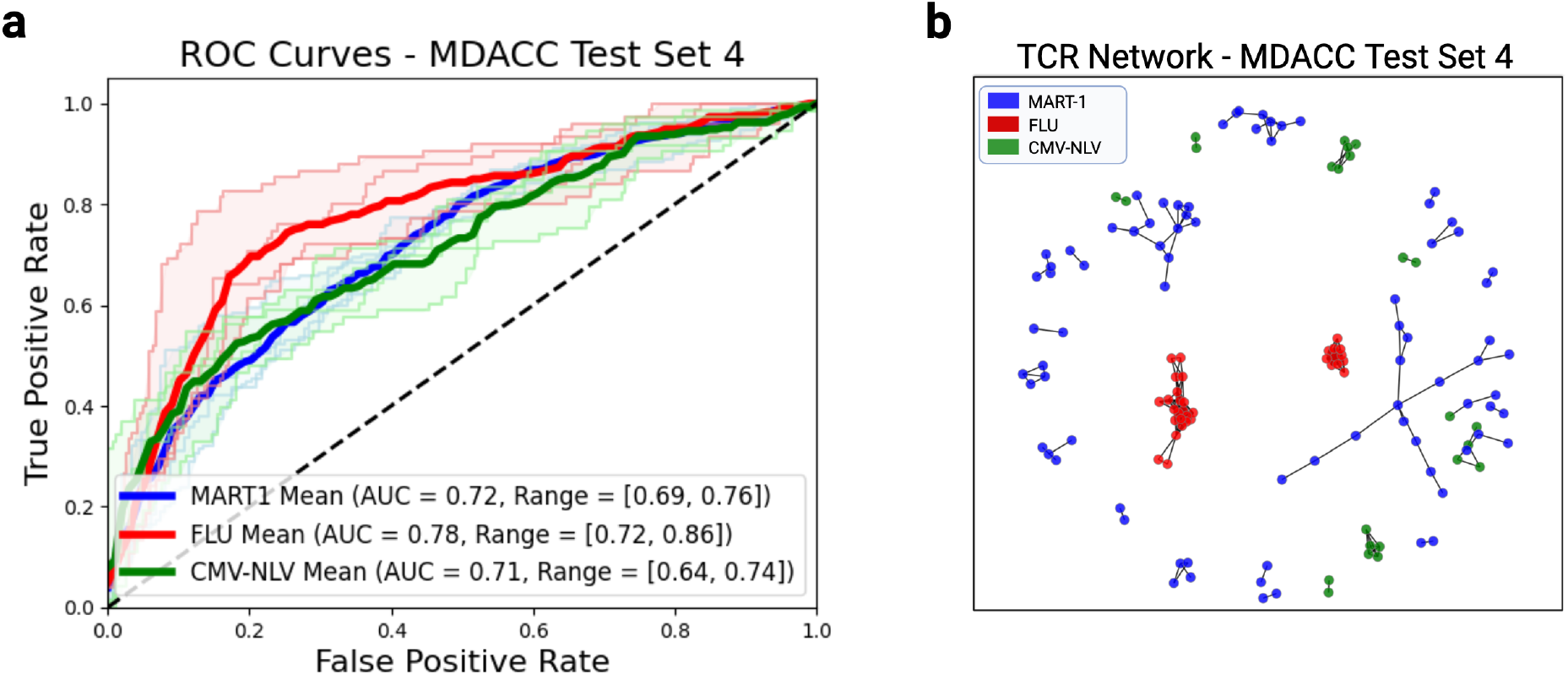
(a) ROC curves for test set 4, which includes *in silico*-derived synthetic 3D structures generated with AlphaFold3 to address the diverse TCR clones for MART-1. The curves illustrate the results for different random subsets excluded from the test set and added to the training set, with their structures generated using AlphaFold3. The average ROC curve is presented alongside the confidence interval (CI) to indicate variability and reliability across random trials. The computed ROC-AUC values for each peptide were reported with 90% confidence intervals, providing a robust assessment of the model’s reliability and predictive power.

To further assess the influence of these generated structures, we compared mutual Q heat maps for the highest- and lowest-performing selections. In the best-performing heat map (Figure S8), for MART-1 structures are learned that the experimental crystal structures form a distinct family, while the *in silico* structures form a separate, non-overlapping group. In contrast, the poorest-performing heat map (Figure S9) shows that, within the same family, the *in silico* and experimental structures share significant structural similarity, indicating that the additional *in silico* structures do not effectively expand the model’s diversity of structural families available for predictions. The experimentally confirmed TCRdist-clustered specificities (Figure 8b) illustrate the relative sequence diversity in *MDACC_5*. In particular, MART-1 exhibits highly diverse TCR clusters, whereas FLU and CMV-NLV display patterns of limited and dense clusters. This result further reinforces the importance of including meaningful structures into the training and optimization steps to appropriately generalize the modeling frameworks to new test data.

Both of the above approaches did not explicitly take into account sequence similarity of test cases relative to existing crystal structures. To evaluate the importance of incorporating new templates that are distinct from existing ones on prediction accuracy, we implemented a third template selection approach (Figure S10a). We first identified clusters that already contained experimental crystal structures (red nodes) and then prioritized TCRs (light blue nodes) from clusters *lacking* existing structures for structure generation. We then repeated the same density-dependent selection of (*n* = 15) ~5% of the *MDACC_5* TCRs to *a priori* confirm their specificity (Figure S10b). The model was then retrained and tested on the remaining data (Figure S10c). This approach led to a slight improvement of ROC-AUC for CMV-NLV, as the relative dearth of available experimental structures for this case provided an opportunity for *in silico*-derived structures to enhance model ability. In contrast, for MART-1 and FLU, the refined selection yielded results similar to those obtained with the (*n* = 30) random selection strategy and the (*n* = 15) representative case selection informed by cluster density and size (Figure 7, 8), indicating that the structural selections were largely consistent across all strategies.

Lastly, to evaluate the robustness of our approach to any particular choice of *in silico* structure creation method, we repeated our analysis again, this time utilizing several alternative approaches, including AlphaFold3 (14), AlphaFold Multimer (30), and Boltz-1 (31) (Figure S11). Repeat predictive assessments demonstrated that overall model performances across these methods were comparable, achieving average ROC-AUC scores for the *MDACC_5* test set of 0.72, 0.78, and 0.71 (AlphaFold3), 0.74, 0.76, and 0.73 (AlphaFold Multimer), and 0.70, 0.76, and 0.72 (Boltz-1) for the Mart1, Flu, and NLV peptides, respectively. These results highlight that our computational approach is robust to any particular method of structure generation, provided that sufficient structural diversity and quality are maintained. Collectively, these results demonstrate the effectiveness of utilizing a small number of structurally informed templates to enhance model generalization to previously unseen TCRs. Additionally, iterative updating with the inclusion of relevant *in silico*-derived structural data can effectively augment the training of the RACER-m model, particularly in scenarios where experimental structural data are limited.

## Discussion

TCR-pMHC prediction is a formidable task, owing both to the diversity of sequence space, together with a sparse sampling of that space in any feasible experimental approach. In this study, we aimed to reliably discern TCR specificity against tumor and viral epitopes in the setting of donor-derived hematopoietic stem cell transplant. Model applications are wide ranging and include, in the context of transplant, identifying tumor antigen-specific TCRs from those that expand during viral reactivation. We thus developed a biophysical modeling framework to appropriately cluster and assign TCR specificity, in addition to an update policy for improving the model as continuous additional patient TCR sequences become available and ultimately validated.

One primary advantage of proceeding with a structure-based approach is the extent to which biophysical interactions can be reliably learned at the level of pairwise TCR-peptide amino acid interactions, which then enable training on a relatively small set of sequences and structures. In this context, patient-derived TCR sequence provide an additional opportunity to refine this approach. Our initial training and predictions benefited from structural clustering to identify antigen-specific TCR clusters corresponding to unique peaks in the distributions of binding interactions, indicating that the RACER-m model performs well in datasets with less diverse TCR repertoires when appropriate structural templates are available. However, test datasets with significant TCR diversity required 1) additional sequence-based clustering and 2) the inclusion of further *in silico*-derived structures for optimal predictions. The first need arose either from an ill-defined mapping of test sequences to multiple template structures or from poor assignment of closely related test sequences to disparate structural templates.

To address these issues, we incorporated additional structural templates generated by AlphaFold3, which expanded the structural diversity within the training set and ensured better coverage of TCR clusters observed in diverse patient datasets. The TCRdist approach effectively captures sequence variations not apparent from structural similarities alone, particularly when templates are limited or TCR diversity is high. Consequently, by integrating these new templates, model predictions became more robust. We remark that the combination of TCRdist assignment and structural model prediction is required for maximal accuracy, where the use of either method to independently assign TCR specificity results in reduced predictive accuracies. These improvements are further supported by the performance of randomly selected training subsets, with the best selection yielding a significantly improved ROC curve due to broader coverage of TCR clusters. Notably, our results suggest that the RACER-m framework maintains predictive utility over a range of structure generation methods, which is reassuring from the standpoint of consistency across structure type.

In our analysis of datasets *MDACC_2, MDACC_3*, and *MDACC_4*, we initially pursued a strategy of identifying a minimal yet representative subset of structural templates capable of capturing TCR specificity. In contrast, the local diversity of *MDACC_5* limited the ability to reliably predict specificity using a fixed set of templates. This finding motivated the inclusion of broader structural coverage to improve model generalizability. While these two strategies contribute opposing effects on the number of available templates, they complement one another meeting the shared objective of selecting structural templates that best represent the distribution of TCRs within each dataset. Despite the above strategy, not all included structures contributed positively to predictive accuracy. Templates exhibiting suboptimal quality or limited representativeness of TCR clusters correlated with poorer performance outcomes. Specifically, we observed an association between poorly performing templates and unfavorable Z-score distributions, suggesting that structural inadequacy—such as inaccurate modeling or poor alignment with real TCR-peptide binding conformations—can distort energetic predictions. These findings emphasize the importance of carefully selecting structural templates to balance coverage and quality, and highlight that both template diversity and accuracy are critical for improving model generalizability across patient cohorts.

Our analysis identified several variations in update procedures whereby new *in silico*-derived structures can be incorporated for more accurate future predictions, which were primarily partitioned based on randomized vs. directed choice of new structural templates selected from diverse sequence clusters. While the strategies explored here offered comparable predictive performance, each case possesses different implications on updating with respect to new information. For example, randomization, while requiring more upfront specificity data, can be used as an unbiased method for incorporating new structures in sequence. In contrast, the selection-based approach, while more efficient in requiring minimal TCR specificity, may result in suboptimal template selection relative to future TCR datasets, particularly if the distribution of sequence diversity becomes skewed in favor of a cluster for future test TCRs. In reality, the best choice is likely context-specific and requires further experimental validation supporting the initial findings herein. We remark that our approach, while achieving high predictive accuracy, still required experimental confirmation of a small sub-sampling of test cases. The principal advantage in applying our structural model is that a few carefully selected templates with sufficient diversity can recover accurate predictions on a majority of unseen test cases.

The improved predictive accuracy of the RACER-m model has important clinical implications for allo-HSCT. By reliably identifying GVL-specific TCRs, clinicians can better select donor-derived TCR repertoires that maximize anti-leukemia effects while minimizing off-target expansions that may lead to graft-versus-host disease (GVHD) or responses to viral antigens. This capability could help refine donor selection and guide therapeutic interventions to improve patient outcomes in allo-HSCT. Future studies could explore expanding the RACER-m framework by incorporating a broader range of peptide targets beyond those studied here. This augmented set could include a larger collection of putative tumor-associated antigens. Additionally, the inclusion of minor-histocompatibility differences (SNP polymorphisms) resulting in differences in self and donor-derived antigens could in the future enable clinicians to add the additional criterion of minimization of GVHD in the optimal selection of a donor TCR repertoire. The integration of sequence-based and structure-based models directly into the training of the energy matrix is one future direction that could provide a more comprehensive approach to capturing the complexity of TCR-pMHC interactions. Another future direction includes leveraging additional molecular features, such as peptide-binding motifs or TCR-peptide docking simulations, to further improve prediction accuracy and extend the applicability of RACER-m to a wider array of antigen-specific immune responses. Our modeling framework in its current form can guide the identification of antigen specific TCRs, and we anticipate that this approach can in the future be expanded to a variety of clinical contexts involving TCR antigen specificity prediction.

## Conflict of Interest Statement

The authors declare that the research was conducted in the absence of any commercial or financial relationships that could be construed as a potential conflict of interest. PJK is a share-holder of Amgen Inc.

## Author Contributions

ZSG, XL, JJM, and JTG designed the study. NT, LZ, XM, HN, DW, YC, QH, SL, and PJK developed the experimental pipeline and performed the experiments. ZSG, XL, and JTG devleoped the computational framework. ZSG performed the computational analysis. ZSG, NT, XL, JJM, and JTG analysed the results. ZSG, XL, JJM, and JTG wrote the first draft. ZSG, NT, LZ, XM, HN, DW, YC, QH, SL, EC, PJK, XL, JJM, and JTG reviewed and edited the manuscript. All authors approve of the final version of the manuscript.

## Acknowledgments

We acknowledge the Tetramer Core Facility at Baylor College of Medicine for providing tetramer reagents used in this study. JTG was supported by the Cancer Prevention Research Institute of Texas (RR210080) and the National Institute of General Medical Sciences of the NIH (R35GM155458). JTG is a CPRIT Scholar in Cancer Research. XL was supported by startup funding from NC State University.

## Data Availability Statement

Sequences and their tetramer and clinical TCR specificities/types will be released on a doi-indexed public repository upon acceptance of the manuscript.

## References

1. Wen Zhang, Peter G Hawkins, Jing He, Namita T Gupta, Jinrui Liu, Gabrielle Choonoo, Se W Jeong, Calvin R Chen, Ankur Dhanik, Myles Dillon, et al. A framework for highly multiplexed dextramer mapping and prediction of t cell receptor sequences to antigen specificity. Science advances, 7(20):eabf5835, 2021.

2. Emma J Grant, Tracy M Josephs, Sophie A Valkenburg, Linda Wooldridge, Margaret Hellard, Jamie Rossjohn, Mandvi Bharadwaj, Katherine Kedzierska, and Stephanie Gras. Lack of heterologous cross-reactivity toward hla-a* 02: 01 restricted viral epitopes is underpinned by distinct αβt cell receptor signatures. Journal of Biological Chemistry, 291(47):24335–24351, 2016.

3. Jason T George, David A Kessler, and Herbert Levine. Effects of thymic selection on t cell recognition of foreign and tumor antigenic peptides. Proceedings of the National Academy of Sciences, 114(38):E7875–E7881, 2017.

4. Kevin Ng Chau, Jason T George, José N Onuchic, Xingcheng Lin, and Herbert Levine. Contact map dependence of a t-cell receptor binding repertoire. Physical Review E, 106(1): 014406, 2022.

5. Barthelemy Meynard-Piganeau, Christoph Feinauer, Martin Weigt, Aleksandra M Walczak, and Thierry Mora. Tulip: A transformer-based unsupervised language model for interacting peptides and t cell receptors that generalizes to unseen epitopes. Proceedings of the National Academy of Sciences, 121(24):e2316401121, 2024.

6. Bjørn PY Kwee, Marius Messemaker, Eric Marcus, Giacomo Oliveira, Wouter Scheper, Catherine J Wu, Jonas Teuwen, and Ton N Schumacher. Stapler: efficient learning of tcr-peptide specificity prediction from full-length tcr-peptide data. bioRxiv, pages 2023–04, 2023.

7. Zahra S Ghoreyshi, Hamid Teimouri, Anatoly B Kolomeisky, and Jason T George. Integration of kinetic data into affinity-based models for improved t cell specificity prediction. Biophysical Journal, 2024.

8. Hamid Teimouri, Zahra S Ghoreyshi, Anatoly B Kolomeisky, and Jason T George. Feature selection enhances peptide binding predictions for tcr-specific interactions. Frontiers in Immunology, 15:1510435.

9. Alessandro Montemurro, Viktoria Schuster, Helle Rus Povlsen, Amalie Kai Bentzen, Vanessa Jurtz, William D Chronister, Austin Crinklaw, Sine R Hadrup, Ole Winther, Bjoern Peters, et al. Nettcr-2.0 enables accurate prediction of tcr-peptide binding by using paired tcrα and β sequence data. Communications biology, 4(1):1060, 2021.

10. Jacob Glanville, Huang Huang, Allison Nau, Olivia Hatton, Lisa E Wagar, Florian Rubelt, Xuhuai Ji, Arnold Han, Sheri M Krams, Christina Pettus, et al. Identifying specificity groups in the t cell receptor repertoire. Nature, 547(7661):94–98, 2017.

11. Zahra S Ghoreyshi and Jason T George. Quantitative approaches for decoding the specificity of the human t cell repertoire. Frontiers in Immunology, 14:1228873, 2023.

12. Xingcheng Lin, Jason T George, Nicholas P Schafer, Kevin Ng Chau, Michael E Birnbaum, Cecilia Clementi, José N Onuchic, and Herbert Levine. Rapid assessment of t-cell receptor specificity of the immune repertoire. Nature Computational Science, 1(5):362–373, 2021.

13. Ailun Wang, Xingcheng Lin, Kevin Ng Chau, José N Onuchic, Herbert Levine, and Jason T George. Racer-m leverages structural features for sparse t cell specificity prediction. Science Advances, 10(20):eadl0161, 2024.

14. Josh Abramson, Jonas Adler, Jack Dunger, Richard Evans, Tim Green, Alexander Pritzel, Olaf Ronneberger, Lindsay Willmore, Andrew J Ballard, Joshua Bambrick, et al. Accurate structure prediction of biomolecular interactions with alphafold 3. Nature, pages 1–3, 2024.

15. Pradyot Dash, Andrew J Fiore-Gartland, Tomer Hertz, George C Wang, Shalini Sharma, Aisha Souquette, Jeremy Chase Crawford, E Bridie Clemens, Thi HO Nguyen, Katherine Kedzierska, et al. Quantifiable predictive features define epitope-specific t cell receptor repertoires. Nature, 547(7661):89–93, 2017.

16. Koshlan Mayer-Blackwell, Stefan Schattgen, Liel Cohen-Lavi, Jeremy C Crawford, Aisha Souquette, Jessica A Gaevert, Tomer Hertz, Paul G Thomas, Philip Bradley, and Andrew Fiore-Gartland. Tcr meta-clonotypes for biomarker discovery with tcrdist3 enabled identification of public, hla-restricted clusters of sars-cov-2 tcrs. Elife, 10:e68605, 2021.

17. Jiawei Zhang, Wang Ma, and Hui Yao. Accurate tcr-pmhc interaction prediction using a bert-based transfer learning method. Briefings in Bioinformatics, 25(1):bbad436, 2024.

18. Shashank Yadav, Dhvani Sandip Vora, Durai Sundar, and Jaspreet Kaur Dhanjal. Tcr-esm: employing protein language embeddings to predict tcr-peptide-mhc binding. Computational and Structural Biotechnology Journal, 23:165–173, 2024.

19. Pieter Moris, Joey De Pauw, Anna Postovskaya, Sofie Gielis, Nicolas De Neuter, Wout Bittremieux, Benson Ogunjimi, Kris Laukens, and Pieter Meysman. Current challenges for unseen-epitope tcr interaction prediction and a new perspective derived from image classification. Briefings in bioinformatics, 22(4):bbaa318, 2021.

20. Anna Weber, Jannis Born, and María Rodriguez Martínez. Titan: T-cell receptor specificity prediction with bimodal attention networks. Bioinformatics, 37(Supplement_1):i237–i244, 2021.

21. David N Garboczi, Deborah T Hung, and Don C Wiley. Hla-a2-peptide complexes: refolding and crystallization of molecules expressed in escherichia coli and complexed with single antigenic peptides. Proceedings of the National Academy of Sciences, 89(8):3429–3433, 1992.

22. Boris Rodenko, Mireille Toebes, Sine Reker Hadrup, Wim JE Van Esch, Annemieke M Molenaar, Ton NM Schumacher, and Huib Ovaa. Generation of peptide–mhc class i complexes through uv-mediated ligand exchange. Nature protocols, 1(3):1120–1132, 2006.

23. Nicole L La Gruta, Stephanie Gras, Stephen R Daley, Paul G Thomas, and Jamie Rossjohn. Understanding the drivers of mhc restriction of t cell receptors. Nature Reviews Immunology, 18(7):467–478, 2018.

24. Tyler Borrman, Jennifer Cimons, Michael Cosiano, Michael Purcaro, Brian G Pierce, Brian M Baker, and Zhiping Weng. Atlas: a database linking binding affinities with structures for wild-type and mutant tcr-pmhc complexes. Proteins: Structure, Function, and Bioinformatics, 85 (5):908–916, 2017.

25. Aram Davtyan, Nicholas P Schafer, Weihua Zheng, Cecilia Clementi, Peter G Wolynes, and Garegin A Papoian. Awsem-md: protein structure prediction using coarse-grained physical potentials and bioinformatically based local structure biasing. The Journal of Physical Chemistry B, 116(29):8494–8503, 2012.

26. Benjamin Webb and Andrej Sali. Comparative protein structure modeling using modeller. Current protocols in bioinformatics, 54(1):5–6, 2016.

27. Mingchen Chen, Xingcheng Lin, Wei Lu, José N Onuchic, and Peter G Wolynes. Protein folding and structure prediction from the ground up ii: Aawsem for α/β proteins. The journal of physical chemistry B, 121(15):3473–3482, 2017.

28. Samuel S Cho, Yaakov Levy, and Peter G Wolynes. P versus q: Structural reaction coordinates capture protein folding on smooth landscapes. Proceedings of the National Academy of Sciences, 103(3):586–591, 2006.

29. Steven Henikoff and Jorja G Henikoff. Amino acid substitution matrices from protein blocks. Proceedings of the National Academy of Sciences, 89(22):10915–10919, 1992.

30. Richard Evans, Michael O’Neill, Alexander Pritzel, Natasha Antropova, Andrew Senior, Tim Green, Augustin Žídek, Russ Bates, Sam Blackwell, Jason Yim, et al. Protein complex prediction with alphafold-multimer. biorxiv, pages 2021–10, 2021.

31. Jeremy Wohlwend, Gabriele Corso, Saro Passaro, Mateo Reveiz, Ken Leidal, Wojtek Swiderski, Tally Portnoi, Itamar Chinn, Jacob Silterra, Tommi Jaakkola, et al. Boltz-1: Democratizing biomolecular interaction modeling. bioRxiv, pages 2024–11, 2024.

32. IEDB. Immune epitope database and analysis resource, 2025. Accessed: 2025-01-19.

