## Supplementary Information for "Biophysical modeling for accurate T cell specificity prediction of viral and tumor antigens"

### S1 Mutual Q

Mutual Q assesses structural similarity between 3D protein configurations, enabling global comparisons of TCR-pMHC structures. This method was used to refine structural templates for MART-1, FLU, and CMV-NLV by identifying and removing templates that exhibited either low structural similarity to others within their group or insufficient resolution. For example, the MART-1-specific structure with PDB ID 5HNT, despite having moderate resolution (3.2 Å), showed poor similarity to other MART-1 TCR-pMHC complexes and was excluded. Similarly, low-resolution structures were identified and removed from the FLU and CMV-NLV groups; for CMV-NLV, one such structure was excluded, leaving two representative templates retained. A full list of removed structures is provided in Table S1, and the resulting clusters are visualized in the Mutual Q similarity heatmap (Figure S1). This refinement process led to the identification of two clusters for MART-1, one for FLU, and two for CMV-NLV, and ensured that only structurally representative templates with sufficient resolution were included for each group.

| Peptides | Removed Outlier Templates (PDB IDs) |
|---|---|
| MART-1 | 5NHT, 6D7G, 5E9D, 3QDJ, 6AMU |
| FLU | 5JHD, 5E6I, 5ISZ, 5EUO |
| CMV-NLV | 5D2L |

Table S1: Outlier templates removed for MART-1, FLU, and CMV-NLV based on mutual Q similarity.

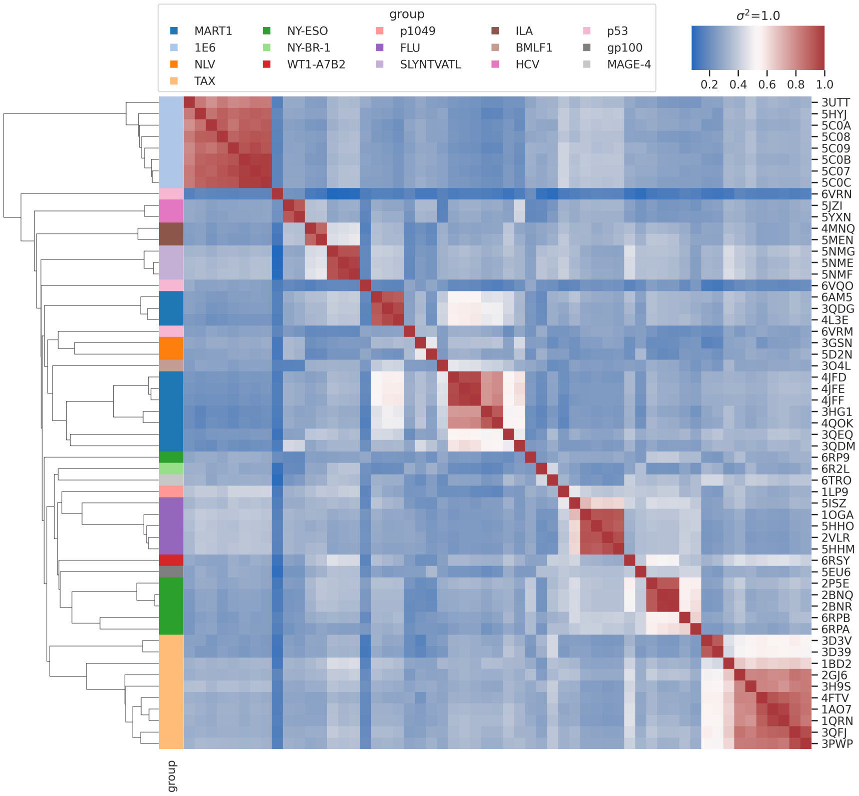


Figure S1: Mutual Q similarity heatmap for refined TCR-pMHC templates. Outlier templates were removed, yielding two MART-1 clusters, one FLU, and two for CMV-NLV clusters (see Table S1)

### S2 TCRdist

#### S2.1 ATLAS Experimental Crystal Structures TCR Network

Our training set comprises both experimental crystal structures from the ATLAS dataset [1] and in silico template structures generated using AlphaFold3 [2]. Figure S2 illustrates the TCR network for experimentally determined crystal structures corresponding to three peptides: MART-1, FLU, and CMV-NLV. This network is plotted based on TCRdist [3, 4], which groups TCRs according to their sequence similarity. To ensure meaningful clustering, we filtered out all clusters containing only a single case. For MART-1 and FLU, this resulted in three and one retained clusters, respectively. In the case of CMV-NLV, all identified clusters consisted of single cases. Therefore, we included two of the three cases and excluded the one with low-resolution data from the CMV-NLV template set. Additionally, Table S2 provides further details on the structural composition of the dataset, listing the PDB IDs for each TCR in the TCR network shown in Figure S2. The table also includes the number of clusters and the mean Z-score distribution for each cluster across the three peptides, offering insights into the structural diversity. Compared to the mutual Q heatmap in Figure S1, the TCR network based on TCRdist exhibits greater structural diversity, reinforcing the need for in silico-generated structures to further enhance training coverage.

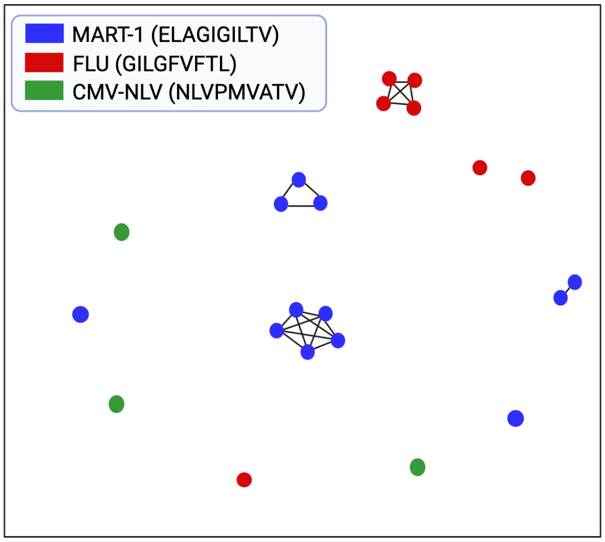


Figure S2: TCR clustering of the ATLAS training set using TCRdist, highlighting sequence-based distances between experimentally identified crystal structure TCRs specific for MART-1, FLU, and NLV epitopes.

| Cluster | Templates | Mean |
|---|---|---|
| MART-1_1 | 3HG1, 3QOK, 4JFD, 4JFF | $\mu_1 = 3.5$ |
| MART-1_2 | 3QDM, 3QEQ | $\mu_2 = 0.9$ |
| MART-1_3 | 3QDG, 3QDJ, 4L3E | $\mu_3 = 6.7$ |
| FLU | 1OGA, 2VLR, 5HHM, 5HHO | $\mu_4 = 4.5$ |
| CMV-NLV_1 | 3GSN | $\mu_5 = 3.9$ |
| CMV-NLV_2 | 5D2N | $\mu_6 = 1.5$ |

Table S2: TCR clusters for MART-1, FLU, and CMV-NLV based on TCRdist similarity, including template structures and mean values, along with the corresponding Z-score means.

### S3 Prediction Prior to Incorporating In Silico Structures and Percentile Calculation

Figures S3 and S4 show ROC curves for *MDACC_3* and *MDACC_4*, respectively, with panel (a) using the Z-score only approach (no *in silico* or percentile), panel (b) employing a percentile-based method with mutual Q refinement, and panel (c) using a percentile-based method with TCRdist refinement. In contrast, Figure S5 (*MDACC_5*) shows panel (a) as Z-score only, panel (b) as percentile only, and panel (c) as the combined percentile and *in silico* approach. Overall, the combined methods (panel (c)) yield the best performance, while the Z-score only approach (panel (a)) performs worst, underscoring the value of incorporating *in silico* structural data.

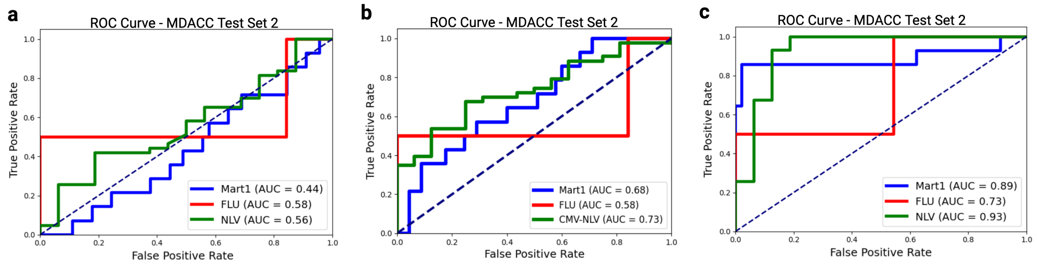


Figure S3: ROC curves for *MDACC_3* comparing: (a) Z-score only (no *in silico* or percentile), (b) percentile-based with mutual Q refinement, and (c) percentile-based with TCRdist refinement.

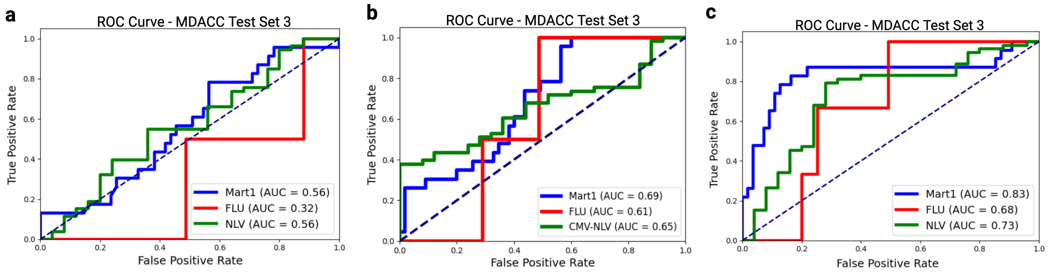


Figure S4: ROC curves for *MDACC_4* comparing: (a) Z-score only (no *in silico* or percentile), (b) percentile-based with mutual Q refinement, and (c) percentile-based with TCRdist refinement.

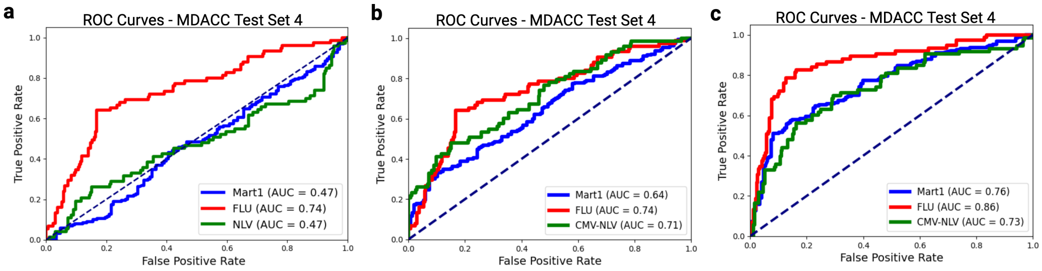


Figure S5: ROC curves for *MDACC_5* comparing: (a) Z-score only (no *in silico* or percentile), (b) percentile only, and (c) percentile-based and *in silico* combined strategy for the most effective random selection

#### S3.1 TCR Network Based on TCRdist

We analyzed the *MDACC_5* dataset (Table 1), which contains TCRs from multiple patients and exhibits a highly diverse range of TCR clusters. TCRdist clustering revealed that this dataset is significantly more diverse than previous patient-derived TCRs, comprising distinct and widely distributed sequences (Figure S2). Given this diversity, we anticipated that existing structural templates alone would be insufficient for accurate predictions and hypothesized that incorporating carefully selected additional templates would enhance model performance.

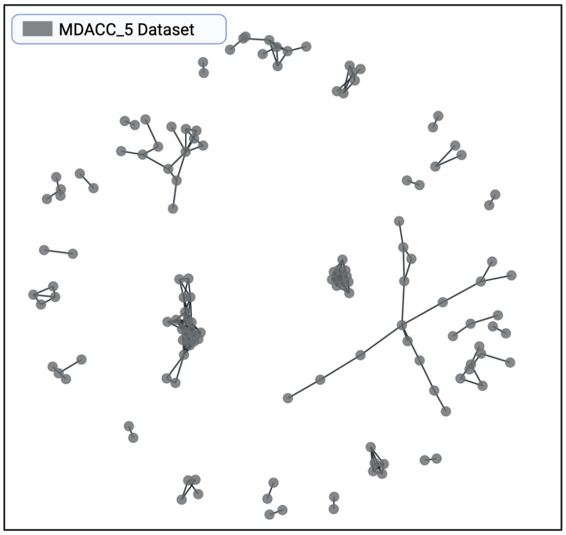


Figure S6: TCRdist clustering of the *MDACC_5* dataset, highlighting its greater structural and sequence diversity compared to previous patient-derived TCRs.

#### S3.2 Random Selection of Thirty Cases for Structure Prediction Using AlphaFold

To address the structural diversity in the *MDACC_5* dataset, we generated *in silico* TCR-pMHC structures using AlphaFold3 [2]. Specifically, we performed six iterations (K=6) of random selection, excluding 30 cases from *MDACC_5* in each iteration. For each selected subset, we generated 3D structures with AlphaFold3 and incorporated them into our existing set of crystal structures (see Results section: Incorporation of *in silico*-derived TCR-pMHC structures generalizes to diverse TCR test sets). The ROC curves for the best- and worst-performing random selections are shown in Figures S7a and S7b, respectively. Figure S7c illustrates the best-performing random selection, where *MDACC_5* TCRs are shown in gray and the selected cases in pink. In contrast, the least effective random selection is depicted in Figure S7d in green, highlighting cases where the added structures were less beneficial. At the time of selection, the peptide specificity of each TCR was unknown; after selection, we retrieved the corresponding peptide data from MDACC and generated 3D structures for the chosen cases before adding them to the training set.

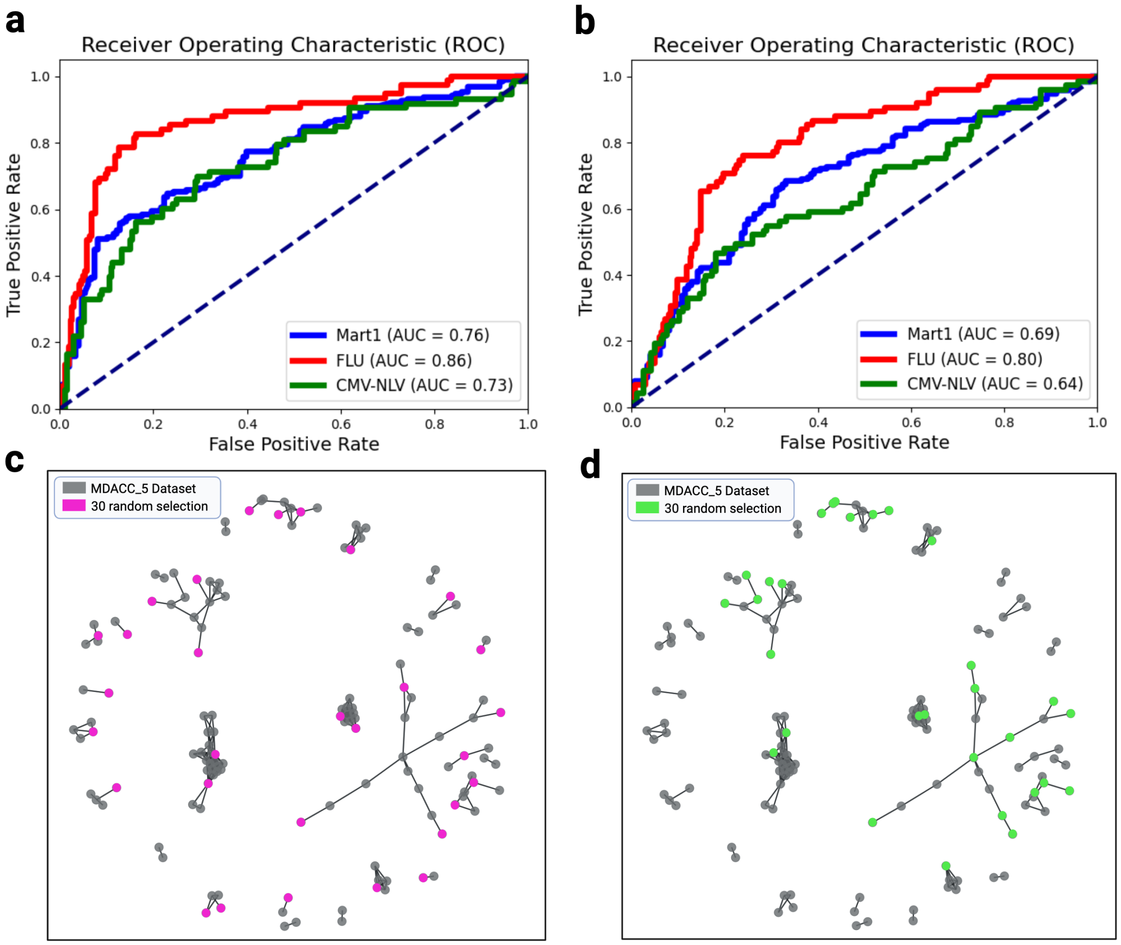


Figure S7: Evaluation of in silico TCR-pMHC structure incorporation into the *MDACC_5* dataset. (a, b) ROC curves for the most and least effective random selections, respectively. (c) TCR clustering of *MDACC_5* (gray) with the most effective selection (pink). (d) TCR clustering with the least effective selection (green).

For the highest-performing selection, we examined the mutual Q heat map (Figure S8) across the MART-1, FLU, and CMV-NLV peptides. In this selection, the heat maps reveal that the experimental crystal structures for MART-1 form a distinct cluster, while the *in silico* structures form a separate, non-overlapping group. Similar segregation is observed for FLU and CMV-NLV, where the additional *in silico* structures provide complementary structural information that is not redundant with the experimental templates. This clear separation of structural families suggests that the selected *in silico* templates introduce unique information, thereby enhancing the diversity of the training set and leading to improved predictive performance.

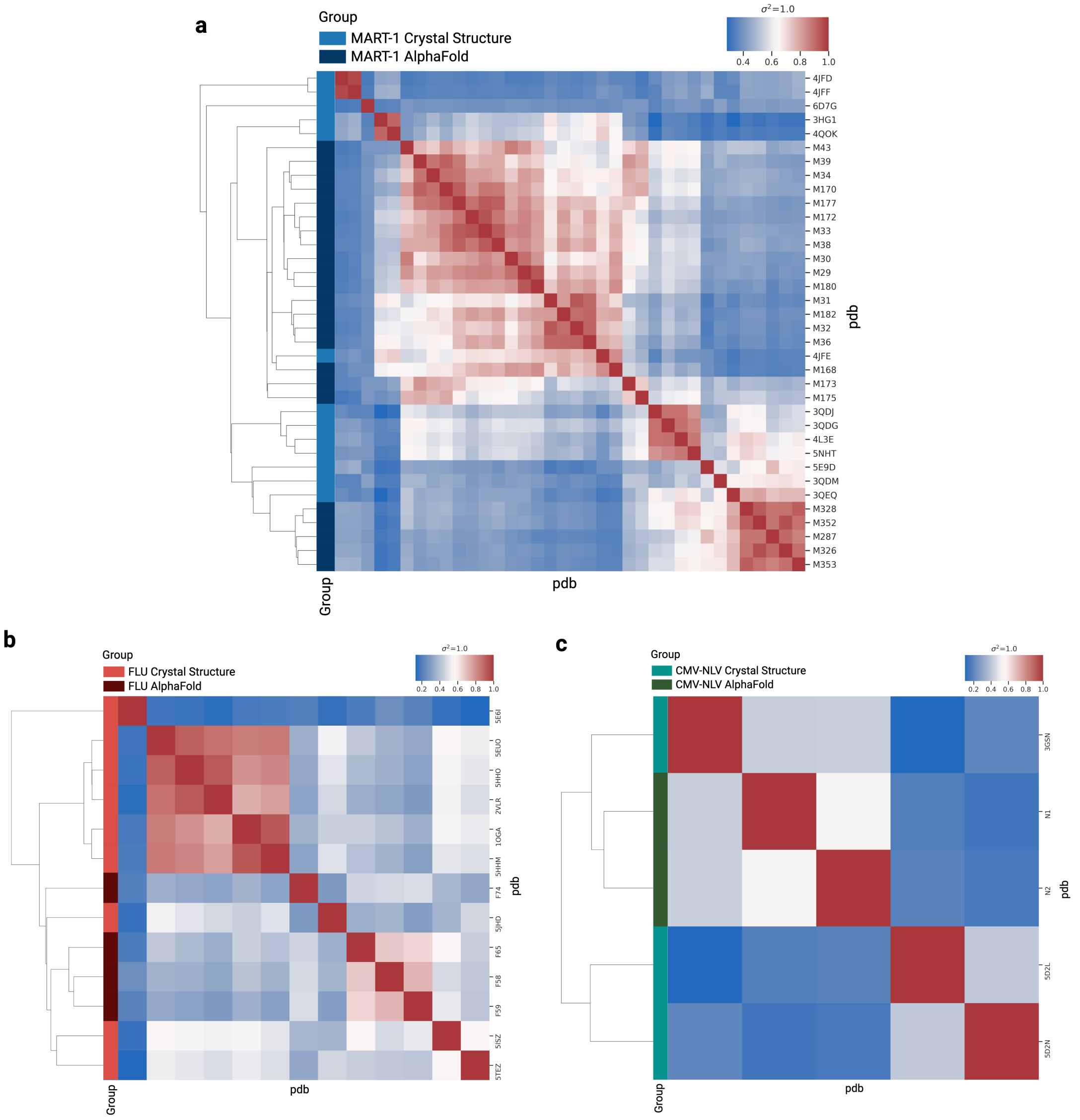


Figure S8: Mutual Q heat maps for the highest-performing selection: (a) MART-1, (b) FLU, and (c) CMV-NLV. For each peptide, experimental crystal structures form a distinct family, while the *in silico*-derived structures segregate into a separate, non-overlapping group, demonstrating that the additional templates contribute novel structural information.

For the lowest-performing selection, we analyzed the mutual Q heat map (Figure S9). In this case, the heat map reveals that both the *in silico* and experimental structures are intermixed within the same structural families, rather than forming distinct groups. This lack of separation indicates that the additional *in silico* structures do not introduce novel structural variations but instead replicate existing families. Consequently, these templates fail to expand the diversity of the structural repertoire, which may contribute to the reduced predictive performance. This analysis underscores the importance of selecting additional templates that offer unique structural information to improve model accuracy.

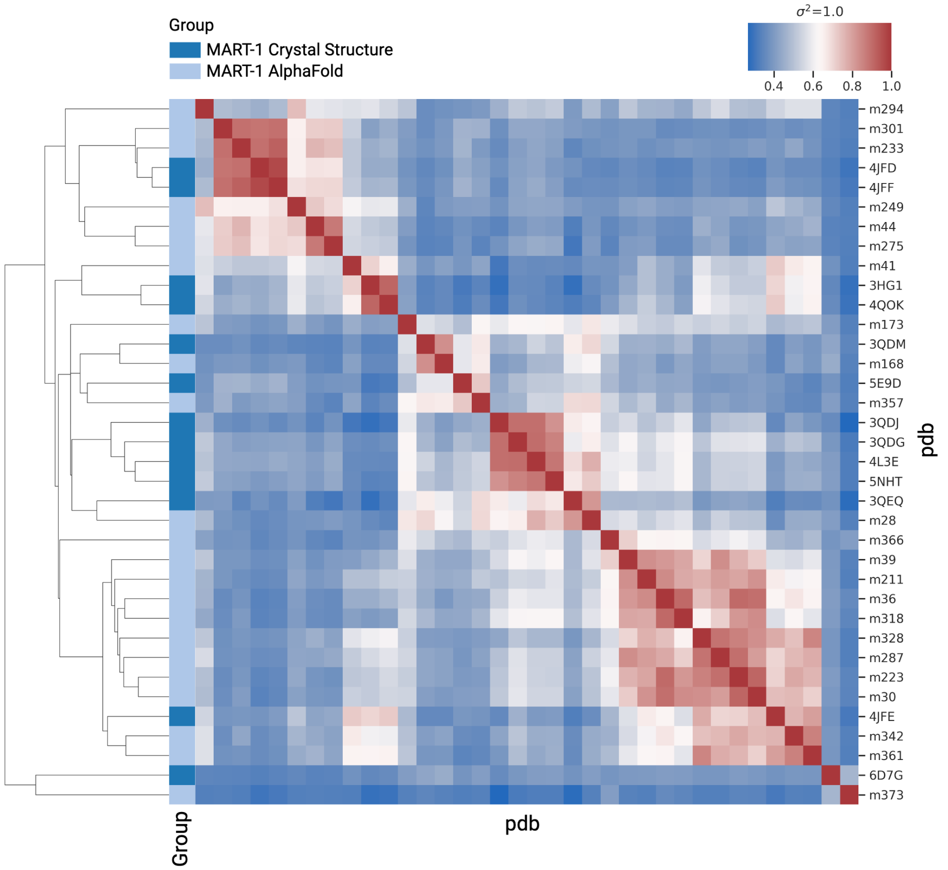


Figure S9: Mutual Q heat map for the lowest-performing selection. Here, both *in silico* and experimental structures co-occur within the same structural families, suggesting that the additional templates do not expand the diversity of structural groups beyond what is captured by existing crystal structures.

#### S3.3 Comparison of Experimental Crystal Structures with TCR Clusters in the *MDACC_5* Dataset

*MDACC_5* is a dataset composed of TCRs from multiple patients, exhibiting a highly diverse range of TCR clusters based on TCRdist [3, 4]. To analyze the clustering patterns, we first applied TCRdist for TCR clustering without prior knowledge of which peptide each TCR recognizes. Additionally, we incorporated the ATLAS dataset [1], which contains experimentally determined crystal structures, to compare its distribution with that of *MDACC_5*. As shown in Figure S10a, the ATLAS dataset appears only in highly dense clusters or as single-case clusters, highlighting that our experimental crystal structures do not sufficiently cover the diverse TCR clusters present in *MDACC_5*. Consequently, many clusters lack at least one corresponding experimental crystal structure, emphasizing the need to generate *in silico* structures to enrich our training set and improve structural coverage.

To address this limitation, we strategically selected 15 TCRs from *MDACC_5* clusters that lacked experimental crystal structures, as illustrated in Figure S10b. Specifically, we first identified clusters containing experimental crystal structures (red nodes) and then prioritized TCRs (light blue nodes) from clusters without experimental structures, ensuring at least one representative from each such cluster was selected for *in silico* structure generation. This approach expanded our structural dataset, bridging gaps where experimental data was unavailable.

The impact of this refined selection strategy is demonstrated in Figure S10c, which presents the ROC curves comparing classification performance. Notably, the ROC-AUC for CMV-NLV improved compared to previous selection strategies. This increase in performance can be attributed to the fact that when no experimental crystal structure is available for a structural family, generating *in silico* structures enhances the model's ability to generalize. In contrast, for structural families that already have representative crystal structures, adding *in silico* structures can introduce noise and reduce performance. For MART-1, we observed higher performance when using the 30 random selections, as additional *in silico* structures were introduced, even for clusters with only two or three cases. However, when using the refined 15-selection strategy, the results remained similar to previous approaches. For FLU, the ROC-AUC values remained comparable across selection strategies, indicating that the refined approach did not significantly alter classification performance for this peptide. Overall, these refinements contributed to a more balanced structural dataset, improving predictive accuracy in cases where experimental structures were initially absent while maintaining consistency where additional structures did not introduce substantial benefits.

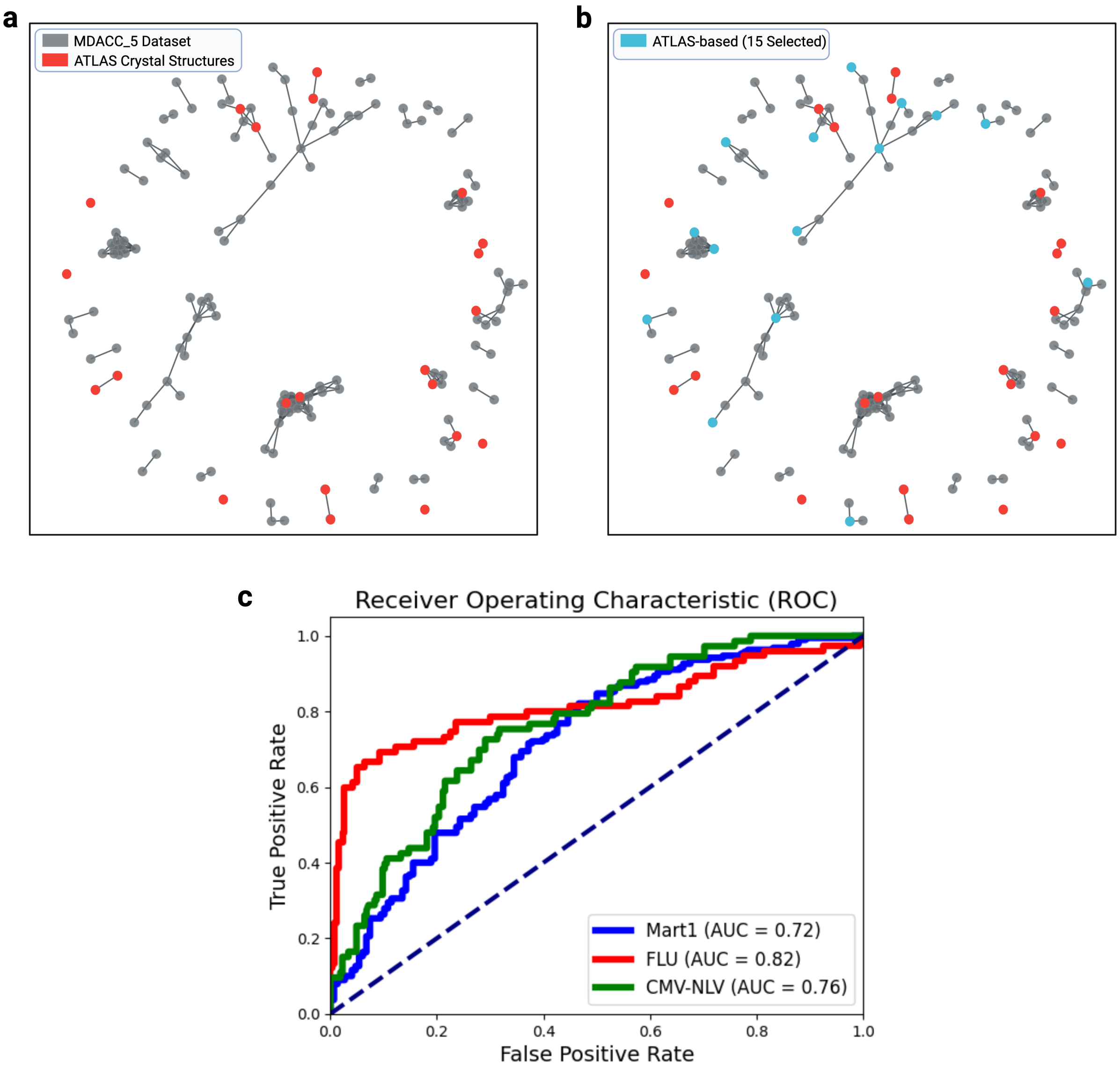


Figure S10: **Comparison of TCR clustering and selection strategies.** (**a**) TCR clustering of the *MDACC_5* dataset based on TCRdist, highlighting ATLAS experimental crystal structures (red), which are predominantly found in dense clusters or as single-case clusters. (**b**) Selection of 15 TCRs (light blue) for *in silico* structure generation from clusters lacking experimental structures. Experimental crystal structures are shown in red, while the remaining *MDACC_5* TCRs are in gray. (**c**)ROC curves comparing the performance of selected cases with the fifteen-case selection based on cluster density.

#### S4 Comparison of In Silico Structure Generation Methods

For comprehensive structural template creation, we utilized three different computational methods: AlphaFold3 [2], AlphaFold Multimer [5], and Boltz-1 [6]. AlphaFold3 employs a diffusion-based architecture to directly predict atomic coordinates, significantly improving prediction accuracy across diverse biomolecular interactions, including proteins, nucleic acids, and small molecules. Its diffusion approach allows for more generalized chemical structure predictions without the extensive use of multiple-sequence alignments (MSAs). AlphaFold Multimer, specifically trained for multi-chain protein complexes, leverages optimized MSA pairing and symmetry handling to enhance prediction accuracy for protein-protein interactions, significantly outperforming previous AlphaFold iterations adapted for multimer prediction. Boltz-1 integrates advancements in data processing and model architecture to broaden access to high-accuracy biomolecular interaction modeling, providing AlphaFold3-level accuracy while enhancing computational efficiency and robustness through improved MSA pairing algorithms and unified cropping methods. Figure S11 illustrates the ROC curves corresponding to Figures 7, 8, S7, and S10, highlighting ROC-AUC scores for MART-1, FLU, and CMV-NLV peptides using these methods and demonstrating their comparable performance and robustness across diverse computational approaches.

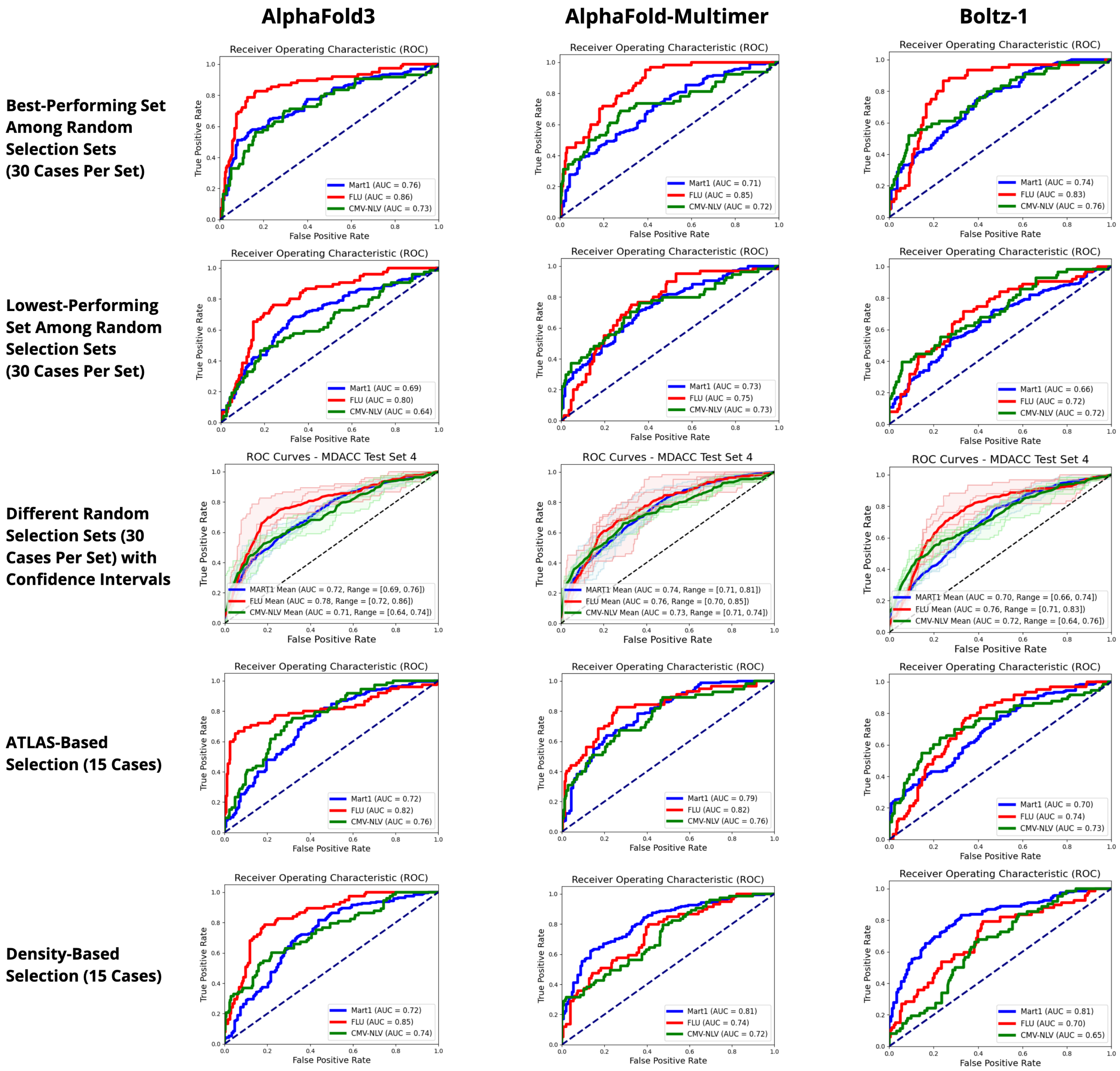


Figure S11: Receiver Operating Characteristic (ROC) curves comparing the predictive performance of AlphaFold3, AlphaFold-Multimer, and Boltz-1 across different structural template selection scenarios. Rows represent various selection methods: Best-performing set among 30 random selections, Lowest-performing set among 30 random selections, 30 random selections with confidence intervals, ATLAS-based selection (15 cases), and Density-based selection (15 cases). Columns represent the three modeling approaches: AlphaFold3, AlphaFold-Multimer, and Boltz-1. ROC curves for Mart1 (blue), Flu (red), and NLV (green) peptides are shown, illustrating the consistency and comparability of prediction accuracies across different methods and selection scenarios.

### References

[1] Tyler Borrman, Jennifer Cimons, Michael Cosiano, Michael Purcaro, Brian G Pierce, Brian M Baker, and Zhiping Weng. Atlas: a database linking binding affinities with structures for wild-type and mutant tcr-pmhc complexes. *Proteins: Structure, Function, and Bioinformatics*, 85(5):908–916, 2017.

[2] Josh Abramson, Jonas Adler, Jack Dunger, Richard Evans, Tim Green, Alexander Pritzel, Olaf Ronneberger, Lindsay Willmore, Andrew J Ballard, Joshua Bambrick, et al. Accurate structure prediction of biomolecular interactions with alphafold 3. *Nature*, pages 1–3, 2024.

[3] Pradyot Dash, Andrew J Fiore-Gartland, Tomer Hertz, George C Wang, Shalini Sharma, Aisha Souquette, Jeremy Chase Crawford, E Bridie Clemens, Thi HO Nguyen, Katherine Kedzierska, et al. Quantifiable predictive features define epitope-specific t cell receptor repertoires. *Nature*, 547(7661):89–93, 2017.

[4] Koshlan Mayer-Blackwell, Stefan Schattgen, Liel Cohen-Lavi, Jeremy C Crawford, Aisha Souquette, Jessica A Gaevert, Tomer Hertz, Paul G Thomas, Philip Bradley, and Andrew Fiore-Gartland. Tcr meta-clonotypes for biomarker discovery with tcrdist3 enabled identification of public, hla-restricted clusters of sars-cov-2 tcrs. *Elife*, 10:e68605, 2021.

[5] Richard Evans, Michael O'Neill, Alexander Pritzel, Natasha Antropova, Andrew Senior, Tim Green, Augustin Žídek, Russ Bates, Sam Blackwell, Jason Yim, et al. Protein complex prediction with alphafold-multimer. *biorxiv*, pages 2021–10, 2021.

[6] Jeremy Wohlwend, Gabriele Corso, Saro Passaro, Mateo Reveiz, Ken Leidal, Wojtek Swiderski, Tally Portnoi, Itamar Chinn, Jacob Silterra, Tommi Jaakkola, et al. Boltz-1: Democratizing biomolecular interaction modeling. *bioRxiv*, pages 2024–11, 2024.
